# *Salmonella* exploits USP32 to coordinate Rab14 and Rab11 recycling pathways for intracellular survival

**DOI:** 10.64898/2026.01.01.696821

**Authors:** Aysegül Sapmaz, Sabina Y. van der Zanden, Jimmy J.L.L Akkermans, Loréline Menager, Lennert Janssen, Esteban François, Natascha Hagoort, Ilana Berlin, Jacques Neefjes, Virginie Stévenin

## Abstract

Many intracellular bacterial pathogens establish membrane-bound niches derived from host material to survive and replicate within host cells. These compartments are carved through subversion of host intracellular trafficking pathways to modulate the composition of the vacuolar membrane. Although recycling-associated small Rab GTPases are frequently observed at infection sites, their role in membrane remodeling and cargo sorting has remained poorly understood. Here, we leverage the extensive early membrane remodeling occurring during *Salmonella* infection to dissect this process. Using endogenously tagged cell lines, we characterize the dynamics of the recycling small GTPases Rab14 and Rab11 at *Salmonella* intracellular infection sites. We report that Rab14 recruits its effector Rufy1, which mediates sorting of CI-M6PR through recycling tubules. Subsequently, Rab11 disperses the newly formed recycling endosomes by interacting with its effector Fip3 for retrograde transport. This pathway is controlled by the deubiquitinating enzyme USP32, which targets both Rab14 and Rab11, enhancing their interactions with Rufy1 and Fip3, respectively. We further show that USP32 is recruited to the infection site via local enrichment of phosphatidylserine-positive membranes, a process triggered by the *Salmonella* effector proteins SopE/E2. Importantly, silencing USP32 significantly impairs intracellular bacterial survival. Our findings reveal a remarkable exploitation of host signaling and trafficking pathways by *Salmonella* to construct a replication-permissive niche. Targeting these pathways may offer new strategies for therapeutic intervention against intracellular bacterial infections.

## INTRODUCTION

Bacterial entry into host cells results in the encapsulation of the internalized pathogen within membrane-bound compartments derived from the plasma membrane. While some bacteria rapidly escape these bacteria-containing vacuoles (BCVs) to access the cytosol, others extensively remodel the limiting membrane to establish a protective and nutrient-rich vacuolar niche. Fusion with endocytic compartments delivers key membrane components that regulate BCV maturation, while selective exclusion of detrimental cargoes promotes bacterial survival.

The facultative intracellular bacterium *Salmonella* provides a striking example of dramatic BCV remodeling within the first hour of infection.^7^ *Salmonella* triggers the formation of *Salmonella*-containing vacuoles (SCVs) within non-phagocytic epithelial cells, by delivering bacterial effector proteins into the host cytosol via a Type-3-Secretion System (T3SS).^1^ Several effectors stimulate actin rearrangement and membrane remodeling, leading to ruffle formation. Ruffle closure next mediates the formation of the SCV alongside infection-associated macropinosomes (IAMs) devoid of bacteria.^2–4^ In approximately 90% of infected epithelial cells, *Salmonella* occupies and replicates within the SCV, pursuing a vacuolar lifestyle. To establish this replication-permissive vacuolar niche, newly formed (*i.e*., nascent) SCVs undergo sequential maturation steps.^5^ While the unique molecular identity of the nascent SCV membrane partly mimics early, sorting, and recycling endosomes,^6^ mature SCVs acquire markers of late endosomes and lysosomes. This transition involves highly dynamic membrane remodeling through fusion with host vesicles and fission of recycling tubules,^7–9^ enabling selective control of the integral SCV membrane protein repertoire. A key feature of this remodeling is the formation of Spacious Vacuole-Associated Tubules (SVATs), which are long, transient tubular structures that emerge from nascent SCVs and mediate both SCV shrinkage and recycling of the cation-independent mannose 6-phosphate receptor (CI-M6PR) away from the SCV membrane.^7,8^ This CI-M6PR removal prevents delivery of lysosomal hydrolases into the SCV lumen, protecting bacteria from degradation.^8,10^

Therefore, the dramatic early remodeling of the nascent SCV membrane is critical for *Salmonella* intracellular pathogenesis. Yet, how *Salmonella* enables this extensive recycling at its infection site is unclear. While host proteins associated with recycling pathways have been observed at *Salmonella*-infection sites via protein overexpression or immunofluorescence,^6,11,12^ their precise function and dynamics during *Salmonella* infection remain poorly characterized. Notably, the small GTPases Rab11 and Rab14, which regulate interconnected recycling pathways, localize near nascent SCVs.^6,11,13^ Both Rabs are targets of the host deubiquitinating enzyme (DUB) USP32,^15^ suggesting a potential regulatory node controlling recycling. Given that post-translational modifications represent common targets exploited by intracellular bacteria to rapidly modulate host protein dynamics,^14^ understanding how the ubiquitination status of recycling machinery is regulated during *Salmonella* infection presents a compelling avenue to decode SCV membrane biogenesis.

Using time-lapse imaging of infected cells expressing endogenously tagged GFP-Rab11 and GFP-Rab14, we reveal a previously unrecognized mechanism controlling Rab GTPase dynamics during *Salmonella* infection. We demonstrate that USP32 promotes multiple Rab-effector interactions involved in recycling and is recruited to infection sites through local enrichment of phosphatidylserine on membranes—a process triggered by the *Salmonella* effectors SopE and SopE2 secreted into the host cytosol upon infection. Importantly, USP32 silencing significantly impairs intracellular bacterial survival. These findings reveal how *Salmonella* exploits host membrane trafficking machinery to establish a replication-permissive vacuolar niche and illuminate pathogen-driven enhancement of host recycling dynamics.

## RESULTS

### Rab14 and Rab11 display distinct and sequential dynamics at *Salmonella* infection sites

SVAT formation requires the Bar-domain protein SNX1, which localizes along those SCV-emerging tubules.^7,8^ To characterize the recruitment of recycling-associated Rabs Rab11 and Rab14 to infection sites, we co-transfected HeLa cells with GFP-SNX1 and either Rab14 or Rab11 overexpression constructs tagged with the far-red fluorophore miRFP670-nano3 (hereafter 670-). Cells were infected with dsRed-expressing *Salmonella enterica* serovar Typhimurium (hereafter *Salmonella*) and imaged by confocal time-lapse microscopy (**Fig. 1**). We observed the recruitment of SNX1 within the same time-frame as *Salmonella* entry suggesting a recruitment within the first minutes of ruffle formation and an expansion of the signal at the infection site along the SVATs (**Fig. 1**). Both Rab11 and Rab14 were recruited after SNX1 at the infection site but displayed distinct sublocalization patterns: Rab14 localized to the limiting membranes of SCVs and IAMs (**Fig. 1A, Video S1**), while Rab11 accumulated in the close vicinity of these compartments (**Fig. 1B, Video S2**). Notably, while Rab14 overexpression did not noticeably affect the reported localization of SNX1 along SVAT,^7,8^ Rab11 overexpression dramatically altered this pattern, with SNX1-positive tubules appearing shorter-lived and less prominent (**Fig. 1A-B, Videos S1-2**).

**Fig. 1:**
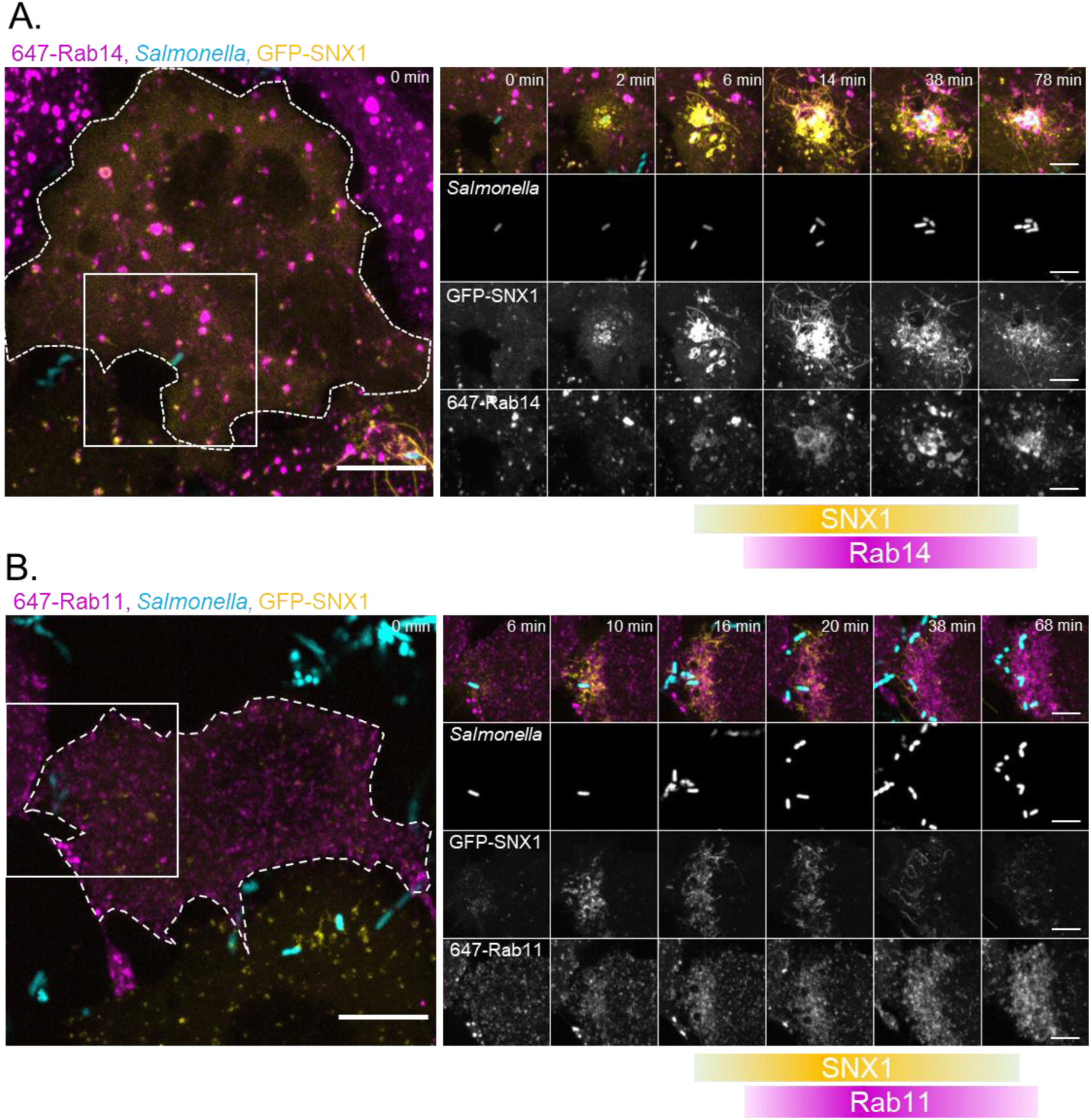
Rab11 and Rab14 are recruited after SNX1 at *Salmonella* infection sites. (A-B) Time-lapse microscopy of HeLa cells overexpressing GFP-SNX1 (yellow), and (A) 670-Rab14 or (B) 670-Rab11 (magenta), infected with dsRed-expressing *Salmonella* (cyan). Bacteria were added to cells immediately before acquisition to capture bacterial entry and early recruitment events. Scale bars: 10 μm (overview) and 5 μm (inset). Representative images from n ≥ 3 biological replicates. See Videos S1–S2.

To characterize Rab14 and Rab11 dynamics at endogenous expression levels, we generated CRISPR-Cas9-edited HeLa cell lines with fluorescently tagged endogenous Rab11 and Rab14 (hereafter eRab11 and eRab14). We validated these cell lines by RT-qPCR (**Fig. S1**) and monitored Rab dynamics during infection without additional overexpression (**Fig. 2A-B, Videos S3-4**). Side-by-side comparison of endogenous Rab11 and Rab14 recruitment revealed divergent localization and dynamics during *Salmonella* infection (**Fig. 2A-B**). eRab14 accumulated at both IAMs and SCVs, displaying uniform membrane distribution, and was transiently observed at SVAT-like tubular extensions emanating from SCVs (**Fig. 2A, Video S3**). The vast majority of SCVs were Rab14-positive (>90 %, **Fig. 2C**), with a median retention time of 30 min at the SCV membrane (**Fig. 2D**). Using fluorescent dextran to label IAMs,^15^ we found that a large fraction of IAMs were Rab14-positive at 30 min post-infection (>40%; **Fig. 2E-F**) but became Rab14-negative within 1 h (**Fig. 2G**). Notably, loss of Rab14 was not accompanied by IAM loss, as these compartments remained dextran-positive after Rab14 depletion, indicating that Rab14 loss reflects maturation of IAM endosomal identity rather than compartment degradation (**Fig. 2F, Video S5**).

**Fig. 2:**
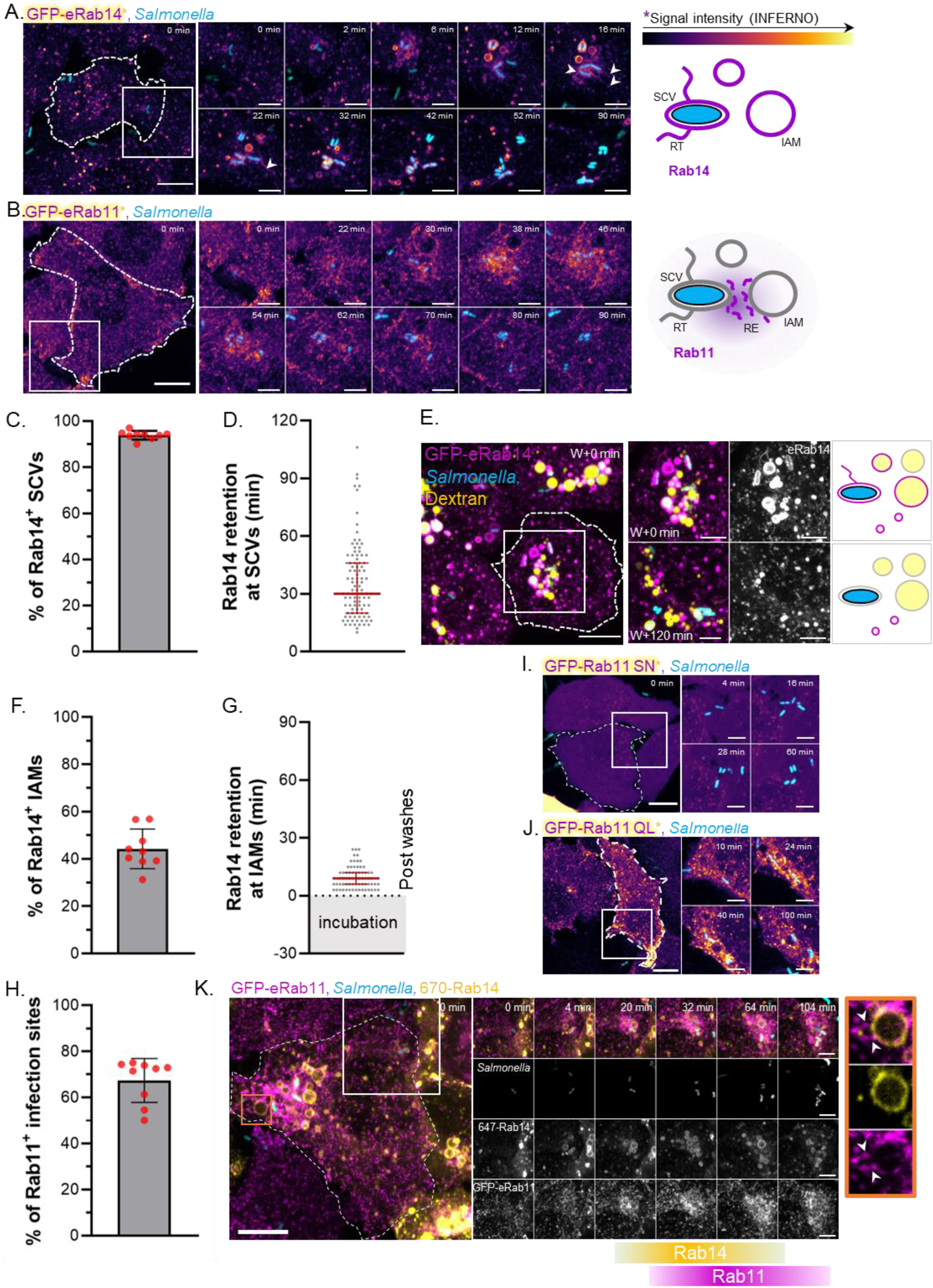
Dynamics of endogenous Rab11 and Rab14 during *Salmonella* infection. (A-B) Time-lapse microscopy of HeLa cells endogenously GFP-tagged for (A) Rab14 or (B) Rab11 (Inferno LUT), infected with dsRed-expressing *Salmonella* (cyan). Bacteria were added immediately before acquisition to capture bacterial entry and early recruitment events. Schematic diagrams depict the interpreted localization of each Rab (magenta) at infection sites. Abbreviations: RT, recycling tubule; SCV, *Salmonella*-containing vacuole; RE, recycling endosome (marked by white arrowhead); IAM, infection-associated macropinosome. Scale bars: 10 μm (overview) and 5 μm (inset). See Videos S3–S4. (C) Percentage of Rab14-positive SCVs during 2 h time-lapse acquisition. Data from three biological replicates; each dot represents one field of view (three fields per replicate). The data represent mean ± SD. (D) Duration of Rab14 retention on SCVs. Each dot represents one SCV from the experiment in C. The data represent median with interquartile range. (E) Time-lapse microscopy of HeLa cells endogenously GFP-tagged for Rab14 (magenta) and infected with dsRed-expressing *Salmonella* (cyan) in the presence of Dextran (yellow). Dextran and bacteria were incubated for 30 min, then washed (W), before acquisition to visualize IAM dynamics. First and last time points are shown. Schematic diagrams depict the interpreted dynamics of Rab14 (magenta) at dextran-labeled IAMs (yellow). See Video S5. (F) Percentage of Rab14-positive IAMs quantified immediately after washing (W + t = 0). Data from three biological replicates; each dot represents one field of view (three fields per replicate). The data represent mean ± SD. (G) Duration of Rab14 retention on IAMs. Each dot represents one IAM from the experiment in F. The data represent mean median with interquartile range. (H) Percentage of Rab11-positive infection sites during 2 h time-lapse acquisition. Data from three biological replicates; each dot represents one field of view (three fields per replicate). The data represent mean ± SD. (I-J) Time-lapse microscopy of HeLa cells overexpressing (I) GFP-Rab11 SN or (J) GFP-Rab11 QL (Inferno LUT), infected with dsRed-expressing *Salmonella* (cyan). Bacteria were added immediately before acquisition to capture bacterial entry and early recruitment events. Scale bars: 10 μm (overview) and 5 μm (inset). (K) Time-lapse microscopy of HeLa cells endogenously GFP-tagged for Rab11 (magenta) and overexpressing 670-Rab14 (yellow), infected with dsRed-expressing *Salmonella* (cyan). Bacteria were added immediately before acquisition to capture bacterial entry and early recruitment events. Scale bars: 10 μm (overview) and 5 μm (inset). White arrowhead, Rab11-positive IAM-emanating tubules. See Video S6.

In contrast to eRab14, eRab11 did not directly coat IAMs and SCVs but instead formed dynamic clouds of small tubulo-vesicular structures in the vicinity of these compartments (**Fig. 2B, Video S4**). This enrichment was observed at most infection sites (>60%, **Fig. 2H**). To confirm that eRab11 enrichment represented membrane-associated pools, we compared recruitment of Rab11 S25N dominant-negative mutant (hereafter Rab11-SN), which locks Rab11 in a GDP-bound inactive state with cytosolic localization, and constitutively active Rab11 Q70L mutants (hereafter Rab11-QL), which locks the GTPase in a GTP-bound active state with membrane-associated localization^16^ (**Fig. 2I-J**). Rab11-SN showed no recruitment to *Salmonella* infection sites, whereas Rab11-QL was enriched in close proximity to SCVs and IAMs, similarly to Rab11 WT (**Fig. 2I-J**), confirming that the observed eRab11 recruitment was membrane-bound. In addition, we observed that Rab11-positive structures did not move toward the infection site but rather appeared *de novo* (**Video S4**), suggesting either the recruitment of Rab11 to pre-existing vesicles at the infection site or the formation of new Rab11-positive structures, potentially through budding from SCVs and IAMs.

To examine the temporal relationship between Rab11 and Rab14 dynamics, we expressed 670-Rab14 in GFP-eRab11 cells and monitored both proteins during *Salmonella* infection. We observed that Rab14 was recruited first to the membranes of SCVs, IAMs, and SVATs, followed by subsequent Rab11 recruitment, which formed a cloud of tubular structures at the periphery of SCVs and IAMs, with some tubules directly budding from these compartments (**Fig. 2K, Video S6**). We confirmed that this recruitment order was not an artifact of overexpression versus endogenous expression by overexpressing 670-Rab11 in GFP-eRab14 cells and observing the same sequence: Rab14 recruitment first at SCVs, IAMs, and SVATs, followed by Rab11 recruitment to nearby structures (**Fig. S2, Video S7**).

Together, detailed time-lapse imaging of endogenously expressed Rab14 and Rab11 reveals their sequential recruitment at distinct membrane localizations of *Salmonella* infection sites, suggesting their coordinated roles in recycling of nascent SCV membrane components.

### The deubiquitinase USP32 targets Rab11 and Rab14 and is required for *Salmonella* **replication**

Rab GTPase dynamics and interactome are highly regulated by posttranslational modifications (PTMs).^16,17^ Notably, recent proteomics screens identified Rab11 and Rab14 as substrates of USP32^18,19^ (**Fig. S3**), a deubiquitinating enzyme (DUB) that regulates intracellular trafficking.^17^ This finding suggests a potential regulatory node controlling both Rabs. Given that bacterial pathogens frequently exploit host PTM systems to rapidly reprogram protein dynamics^14^ and the therapeutic appeal of targeting deubiquitinases,^20^ we investigated whether USP32 regulates Rab14 and Rab11 dynamics during *Salmonella* infection.

We first validated that USP32 deubiquitinates Rab14 and Rab11 using *in vitro* ubiquitination assays. For each Rab, we co-expressed GFP-Rab, HA-ubiquitin, and either myc-USP32 WT or the catalytically inactive myc-USP32 C743A mutant, with vector-only controls as reference (**Fig. 3A-D**). Expression of USP32 WT, but not the C743A mutant, significantly reduced ubiquitination of both Rab14 (**Fig. 3A-B**) and Rab11 (**Fig. 3C-D**). These results confirm that USP32 deubiquitinase activity removes ubiquitin from these substrates.

**Fig. 3:**
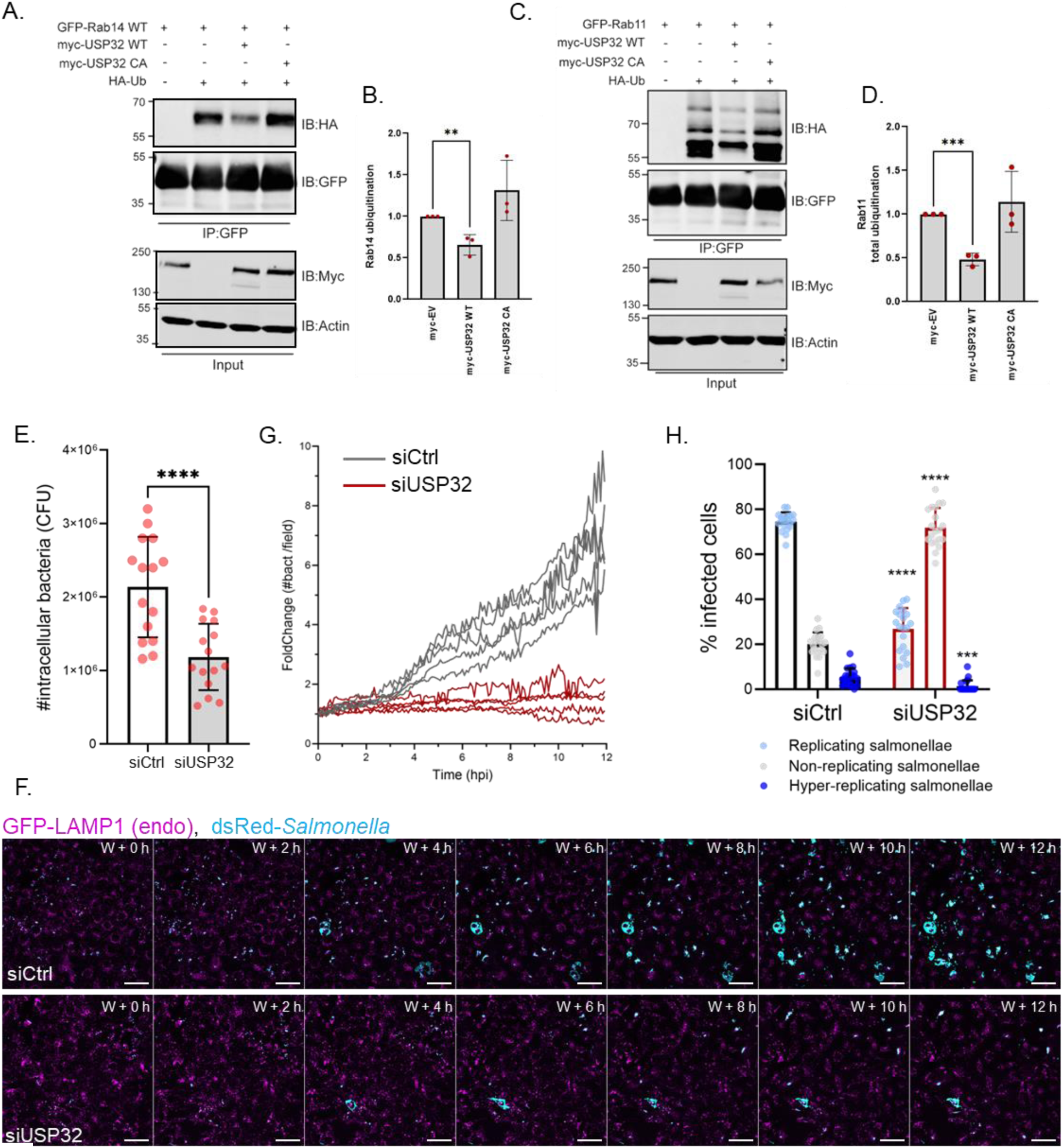
USP32 deubiquitinates Rab11 and Rab14 and is required for *Salmonella* replication. (A-B) Representative immunoblot (A) and quantification (B) of GFP-Rab14 ubiquitination levels in HEK293T cells co-expressing HA-ubiquitin with wild-type USP32, catalytically inactive USP32 C743A, or empty vector control. GFP-Rab14 was immunoprecipitated and ubiquitination was detected by anti-HA immunoblot (n=3 biological replicates). The data represent mean ± SD. Statistical significance was determined by ordinary one-way ANOVA. (C-D) Representative immunoblot (C) and quantification (D) of Rab11 ubiquitination under the same experimental conditions as in A-B (n=3 biological replicates). (E) Quantification of intracellular *Salmonella* at 18 hpi by gentamicin protection assay in siCtrl versus siUSP32 conditions (n=5 biological replicates, each performed in technical triplicate). The data represent mean ± SD. Statistical significance was determined by *t*-test. (F) Representative time-lapse images of HeLa cells with endogenously GFP-tagged LAMP1 (magenta) infected with dsRed-*Salmonella* (cyan) under siCtrl versus siUSP32 conditions (n>3 biological replicates). Acquisition began immediately after 30 min incubation with bacteria and washing (t = W + 0). See Video S8. (G) Quantification of bacterial count per field of view over 12 h under siCtrl versus siUSP32 conditions (n = 5 fields per condition; representative of 3 biological replicates). (H) Single-cell analysis of *Salmonella* replication dynamics under siCtrl versus siUSP32 conditions, based on time-lapse microscopy (n = 4 biological replicates with 5 fields per condition and replicate). The data represent mean ± SD. Statistical significance was determined by *t*-test performed between siCtrl and siUSP32 conditions for each category of replications.

We next assessed the impact of USP32 silencing on *Salmonella* intracellular survival and replication. A gentamicin protection assay revealed a significant reduction in intracellular bacterial numbers at 18 h post-infection (hpi) of cells with depleted USP32 expression (**Fig. 3E**). To account for heterogeneity in *Salmonella* replication dynamics that could confound bulk measurements, we performed single-cell time-lapse microscopy of *Salmonella*-infected HeLa cells endogenously expressing GFP-LAMP1 (labeling the SCV) in siCtrl and siUSP32 conditions (**Fig. 3F-G**). *Salmonella* replication in USP32-depleted cells was mostly abrogated, beginning from the first hours of infection (**Fig. 3G**). Quantification of single-cell replication dynamics revealed a significant increase in non-replicating intracellular *Salmonella*, indicating that in the absence of USP32, SCVs become non-permissive for bacterial replication (**Fig. 3H**).

### USP32 modulates the Rab14 interactome to regulate effector recruitment

USP32 has previously been shown to modulate the interaction preferences of its protein targets Rab7 and LAMTOR, either enhancing or decreasing their affinity for specific binding partners.^18,19^ To determine whether USP32 similarly impacts the Rab14 interactome, we examined Rab14’s affinity for key interactors upon USP32 silencing, including the Rab14 effectors Nischarin and Rufy1, as well as the Rab14 GAP AS160.

The Rab14 effector Nischarin has been reported to localize at *Salmonella* infection sites together with Rab14.^13^ Co-immunoprecipitation experiments revealed that USP32 silencing resulted in decreased interaction between Rab14 and Nischarin (**Fig. 4A-B**). Although Nischarin has been shown to increase the membrane residency of Rab14,^13^ we did not observe changes in Rab14 dynamic recruitment at infection sites upon USP32 silencing (**Fig. S4**). We therefore hypothesized that USP32 might also regulate other factors controlling Rab14 membrane association. We examined AS160, a GAP that promotes Rab14 GTP hydrolysis and thus its membrane removal. Co-immunoprecipitation assays demonstrated a significant increase in Rab14-AS160 interactions upon USP32 silencing (**Fig. 4C-D**). In addition, Nischarin silencing also significantly increased AS160 binding to Rab14 (**Fig. 4C-D, Fig. S5A**), validating the previously proposed hypothesis that Nischarin may prevent this interaction.^13^ These findings suggest that USP32 simultaneously decreases Rab14 interactions with both AS160 and Nischarin, resulting in antagonistic effects on Rab14 membrane residency that may neutralize each other. This model is consistent with the lack of observable impact of USP32 on Rab14 dynamics during *Salmonella* infection (**Fig. S4**).

**Fig. 4.**
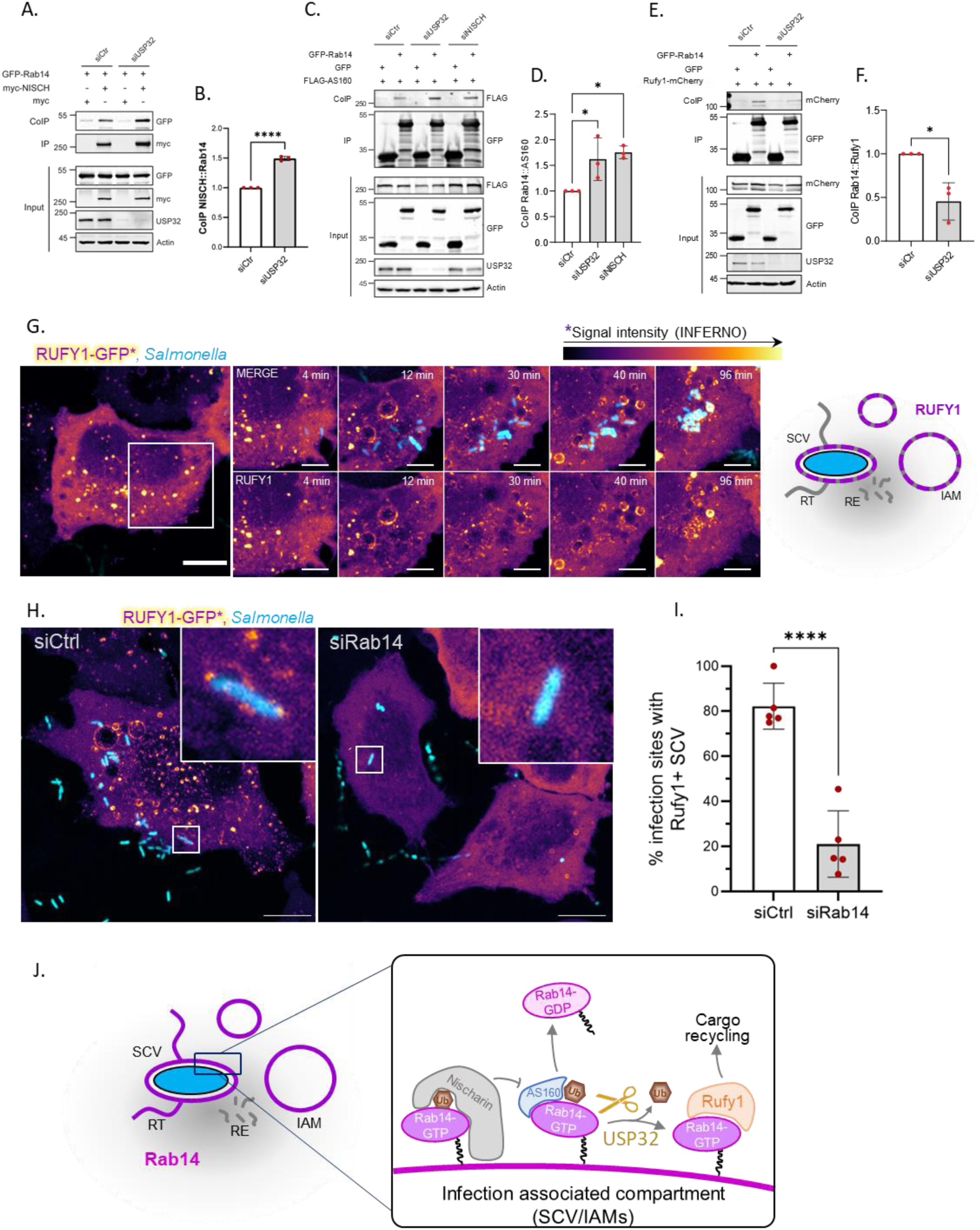
Rab14 interactions are shifted by on USP32. (A-B) Representative immunoblot (A) and quantification (B) of co-immunoprecipitation (CoIP) of myc-Nischarin (NISCH) with GFP-Rab14 in siCtrl versus siUSP32 conditions (n=3 biological replicates). The data represent mean ± SD. Statistical significance was determined by unpaired *t*-test. (C-D) Representative immunoblot (C) and quantification (D) of co-immunoprecipitation (CoIP) of FLAG-AS160 with GFP-Rab14 in siCtrl, siUSP32 and siNischarin (siNISCH) conditions (n=3 biological replicates). Nischarin silencing was validated by RT-qPCR (Fig. S5A). The data represent mean ± SD. Statistical significance was determined by ordinary one-way ANOVA. (E-F) Representative immunoblot (E) and quantification (F) of the co-immunoprecipitation (CoIP) of Rufy1-mCherry with GFP-Rab14 in siCtrl versus siUSP32 conditions (n=3 biological replicates). The data represent mean ± SD. Statistical significance was determined by unpaired *t*-test. (G) Time-lapse microscopy of HeLa cells overexpressing Rufy1-GFP (Inferno LUT), infected with dsRed-expressing *Salmonella* (cyan). Bacteria were added immediately before acquisition to capture bacterial entry and early recruitment events. Schematic diagrams depict the interpreted localization of Rufy1 (magenta) at infection sites. Abbreviations: RT, recycling tubule; SCV, *Salmonella*-containing vacuole; RE, recycling endosome; IAM, infection-associated macropinosome. Top panel, merge; bottom panel, Rufy1-GFP alone. Scale bars: 10 μm (overview) and 5 μm (inset). See Video S9. (H-I) Representative microscopy acquisition (H) and quantification (I) of GFP-Rufy1 recruitment at the SCV in siCtrl versus siUSP32 conditions (n=5 time-lapse acquisition, > 75 quantified cells per condition). The data represent mean ± SD. Statistical significance was determined by *t*-test. (J) Model of USP32 regulation of the Rab14 interactome. Ubiquitinated Rab14 preferentially interacts with Nischarin and AS160. Nischarin stabilizes Rab14 at the membrane by blocking Rab14-AS160 interaction, whereas AS160 promotes GTP hydrolysis and Rab14 release from the membrane. USP32-mediated deubiquitination of Rab14 shifts its binding preference toward the effector Rufy1, which mediates membrane cargo sorting. Through this mechanism, USP32 promotes Rab14-dependent cargo recycling functions.

The Rab14 effector Rufy1 is a member of the RUFY (RUN and FYVE domain-containing proteins) protein family^21^ involved in SNX1-mediated sorting of CI-M6PR from endosomes to the trans-Golgi network.^22^ *Salmonella* can prevent the accumulation of CI-M6PR from the SCV by perturbing its trafficking.^10^ In addition, we observed the presence of M6PR on some SVATs at 1 hpi (**Fig. S6**), suggesting that *Salmonella* not only prevents M6PR trafficking towards the SCV but also sorts the CI-M6PR present at its limiting membrane, consistent with the previous observation that SNX1 silencing resulted in an accumulation of CI-M6PR at *Salmonella* infection sites.^8^ As Rufy1 is a known actor of CI-M6PR sorting through SNX1-positive tubules,^22^ we investigated the USP32-dependent affinity of Rab14 for Rufy1. Co-immunoprecipitation experiments revealed a significant decrease in Rufy1-Rab14 interaction upon USP32 silencing (**Fig. 4E-F**). To explore the relevance of this interaction during *Salmonella* infection, we performed time-lapse imaging of *Salmonella*-infected cells overexpressing GFP-Rufy1. We observed the transient recruitment of Rufy1 to the limiting membrane of SCVs and IAMs in a punctate pattern, suggesting the recruitment of Rufy1 to a specific membrane domain (**Fig. 4G, Video S9**). In addition, Rufy1 was observed at the SCV after Rab14 recruitment (**Fig. S7**), and Rufy1 recruitment to SCVs was drastically reduced upon Rab14 silencing (**Fig. 4H-I, Fig. S5B**). Together, these data suggest that USP32 shifts the protein interaction preferences of Rab14, promoting its interaction with Rufy1, a known mediator of membrane cargo recycling^22^ (**Fig. 4J**). These results are consistent with earlier observations of CI-M6PR recycling alteration upon USP32 silencing.^18^

### USP32 modulates Rab11-FIP3 interaction and regulates Rab11 dynamics at the infection site

To assess the effect of USP32 on Rab11 dynamics at the infection site, we performed time-lapse imaging of *Salmonella*-infected GFP-eRab11 HeLa cells treated with siUSP32 or siCtrl. USP32 silencing significantly increased the maximal amplitude of Rab11 recruitment to the infection site (**Fig. 5A-B**). Strikingly, siUSP32-treated cells also displayed Rab11 accumulation at a perinuclear location, likely corresponding to the centrosome, suggesting altered Rab11 interaction with motor proteins in the absence of USP32 (**Fig. S8**). The interaction of Rab11 with cytoskeletal transport is regulated by the Rab11FIP protein family (hereafter FIP).^23^ Among these, FIP3 links Rab11 to dynein for retrograde transport.^24^ We analyzed FIP3 localization during infection through time-lapse microscopy of *Salmonella*-infected HeLa cells overexpressing FIP3. We observed transient FIP3 recruitment to the infection site, localizing at a vesicle cloud similar to that observed for Rab11 (**Fig. 5C**). These FIP3-positive structures exhibited retrograde movement away from the infection site (**Video S10**).

**Fig. 5.**
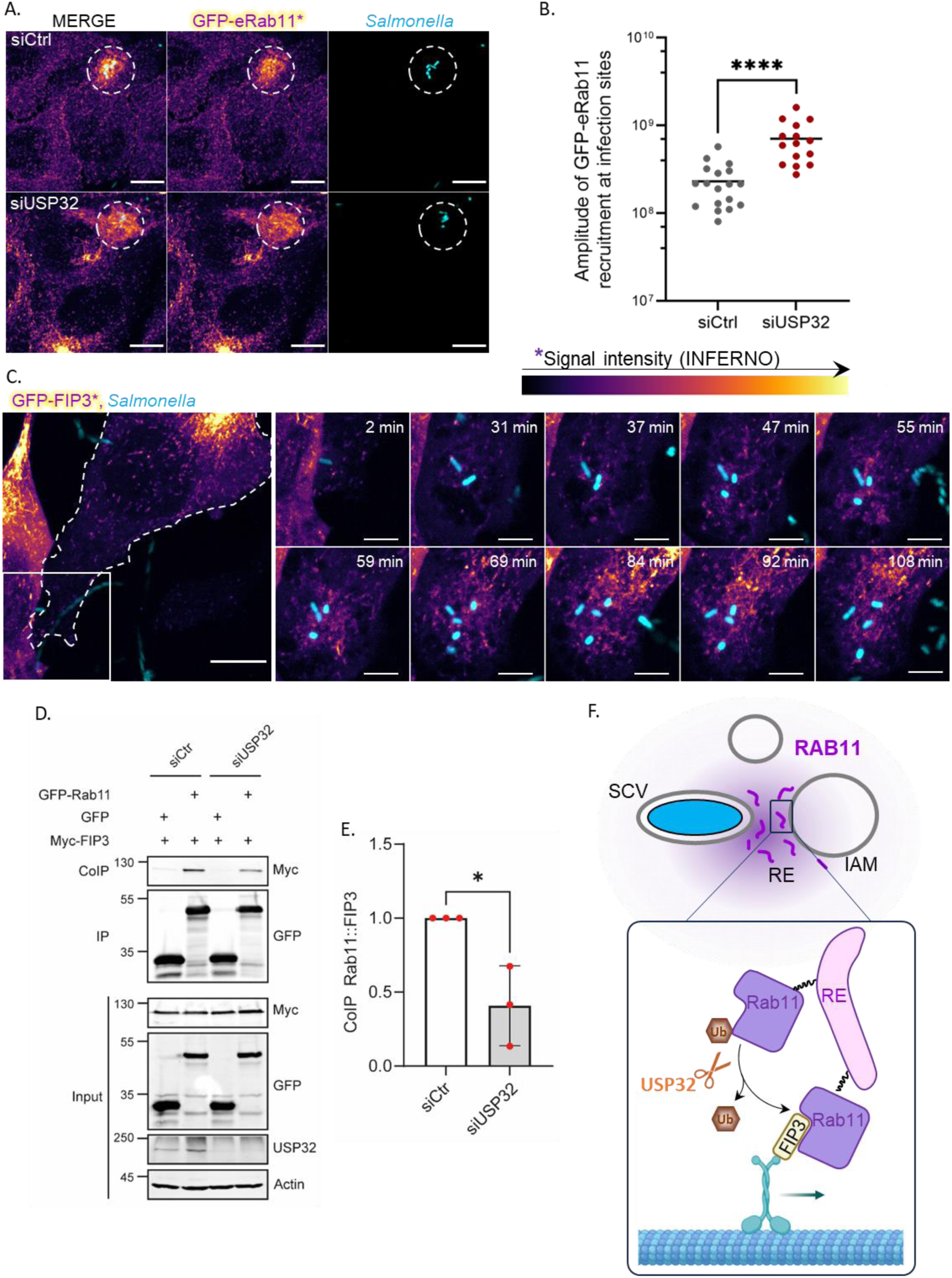
USP32 promotes Rab11–FIP3 interactions and regulates Rab11 dynamics at the infection site. (A-B) Representative microscopy image (A) and quantification (B) of GFP-eRab11 recruitment intensity at *Salmonella* infection sites, quantified by time-lapse imaging of HeLa cells endogenously expressing GFP-Rab11 and infected with *Salmonella*, in siCtrl versus siUSP32 conditions. Each point represents one infection site. Statistical significance was determined by *t*-test. (C) Time-lapse microscopy of HeLa cells overexpressing GFP-FIP3 (Inferno LUT), infected with dsRed-expressing *Salmonella* (cyan). Bacteria were added immediately before acquisition to capture bacterial entry and early recruitment events. Scale bars: 10 μm (overview) and 5 μm (inset). See Video S10. (D-E) Representative immunoblot (D) and quantification (E) of co-immunoprecipitation (CoIP) of Myc-FIP3 with GFP-Rab11 in siCtrl versus siUSP32 conditions (n=3 biological replicates). The data represent mean ± SD. Statistical significance was determined by unpaired *t*-test. (F) Model of USP32-mediated regulation of Rab11 dynamics. USP32 deubiquitinates Rab11, promoting its binding to the effector FIP3, which mediates retrograde transport. Through this mechanism, USP32 promotes the evacuation of recycled cargo from the infection site.

We next examined whether USP32 regulates the FIP3-Rab11 interaction. Co-immunoprecipitation experiments revealed that USP32 silencing significantly decreased FIP3-Rab11 binding, demonstrating that USP32 promotes this interaction (**Fig. 5D-E**). Together, these findings suggest that USP32, by enhancing the FIP3-Rab11 interaction, facilitates efficient retrograde displacement of recycling endosomes and prevents their accumulation near the SCV at the infection site (**Fig. 5F**).

### Phosphatidylserine accumulation at the infection site promotes USP32 recruitment

Given that the *Salmonella* infection site exhibits intense recycling dynamics that could be promoted by USP32, we investigated whether USP32 localization contributes to this process. GFP-USP32 displayed predominantly cytosolic localization in HeLa cells, whereas its catalytically inactive mutant GFP-USP32 C743A showed increased membrane association (**Fig. S9**). This pattern suggests that USP32 resides primarily in the cytosol and is transiently localized at membranes during catalytic activity, making direct observation of its recruitment dynamics to infection sites challenging.

To overcome this limitation and account for the fact that *Salmonella* infection triggers alterations in membrane phospholipid composition,^1,2^ we analyzed the lipid-binding properties of recombinant USP32 using a PIP strip assay. Remarkably, USP32 preferentially bound phosphatidylserine (PS) and phosphatidic acid (PA) (**Fig. 6A**). To validate this finding in cells, we examined the colocalization of USP32 with PS using the PS biosensor LactC2.^25^ Since PS is primarily restricted to the inner leaflet of the plasma membrane, we performed immunofluorescence imaging of cells co-expressing GFP-USP32 and RFP-LactC2, followed by anti-GFP immunostaining to enhance visualization of the membrane-bound fraction of USP32 (**Fig. 6B**). We observed colocalization of USP32 and LactC2 in subcortical regions, in line with the *in vitro* lipid-binding specificity of USP32 for PS.

**Fig. 6.**
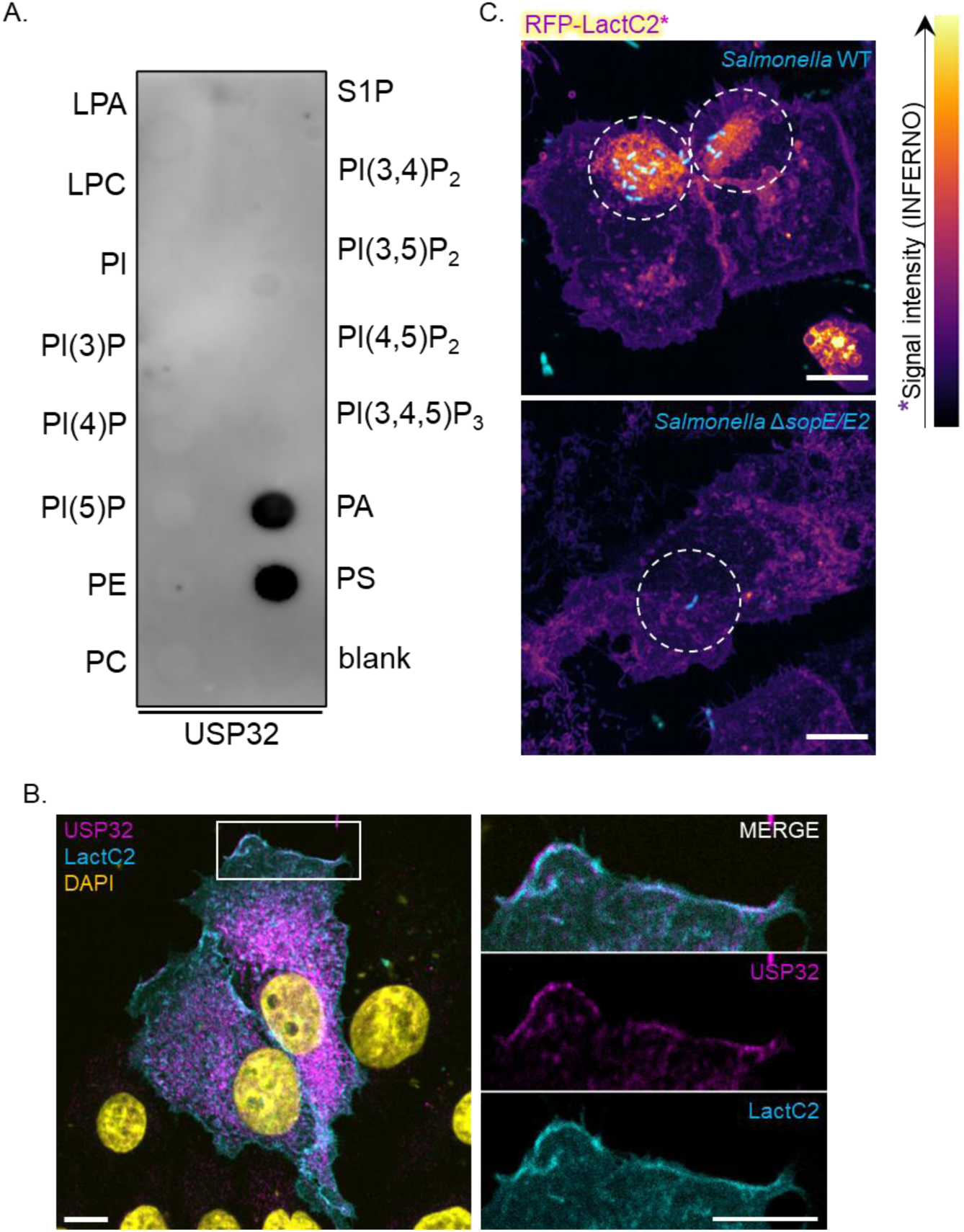
Local changes of lipid composition at the infection site favor USP32 recruitment. (A) Pip strip assay of recombinant USP32, representative of three biological replicates. (B) Colocalization of USP32 with phosphatidylserine (PS). HeLa cells expressing GFP-USP32 and the PS biosensor RFP-LactC2 (cyan) were immunostained with anti-GFP antibody (magenta). Scale bar: 10 µm. (C) Live cell imaging of RFP-LactC2 expressing cells (PS biosensor, Inferno LUT), infected by WT (top panel) or Δ*sopE/E2* (bottom panel) *Salmonella* (cyan). Scale bar: 10 µm.

Given the extensive formation of plasma membrane-derived compartments during *Salmonella* infection through membrane ruffling and IAM formation, we hypothesized that these processes could drive PS accumulation at the infection site. Time-lapse microscopy of cells expressing the LactC2 PS biosensor revealed a striking increase in PS at WT *Salmonella* infection sites (**Fig. 6C, Video S10**). Membrane ruffle formation is triggered by the coordinated action of multiple *Salmonella* effectors.^1^ Specifically, SopE and SopE2 drive actin cytoskeleton remodeling required for ruffle and IAM formation.^2^ We observed that infection with a *Salmonella* Δ*sopE/E2* double mutant abolished PS accumulation at the infection site, demonstrating that SopE/E2-dependent membrane remodeling is essential for this process.

Together, these results reveal a novel function for IAMs as platforms that concentrate PS at the infection site, thereby recruiting PS-binding host proteins such as USP32 to promote recycling dynamics during infection.

## DISCUSSION

Selective membrane cargo sorting mechanisms are subjects of intense investigation due to their fundamental roles in protein homeostasis and cellular signaling.^26^ However, these processes remain less well characterized in host-pathogen interactions. Intracellular pathogens can serve as powerful tools to enhance and dissect such cellular processes. To survive and replicate within host cells, intracellular pathogens actively remodel the intracellular environment to establish favorable conditions. The deep rewiring of host signaling pathways can even promote cell transformation and cancer development, showcasing the capacity of bacteria to exacerbate cell biological processes.^27,28^ A classic example is provided by *Salmonella*, which invades, manipulates host biochemistry, and extensively remodels the endocytic system of intestinal epithelial cells to establish its intracellular replication niche. By decorating its nascent SCV with a hybrid early/recycling endosomal identity, *Salmonella* enables simultaneous maturation of its vacuole while eliminating bactericidal proteins from the SCV limiting membrane.^7^

Two decades ago, Smith and colleagues demonstrated that SCV maturation was impaired when recycling was blocked, proposing that “protein movement through the endocytic recycling system is regulated through at least two concurrent pathways and that efficient interaction with these pathways is necessary for maturation of the *Salmonella*-containing vacuole.”^11^ 20 years later, we here identified several recycling pathways that operate in concert under spatiotemporally regulated PTM control to orchestrate the complex maturation dynamics of the SCV.

Small Rab GTPases are key orchestrators of membrane trafficking dynamics. Although regularly detected at infection sites, the mechanisms by which bacteria manipulate Rab proteins associated with recycling have remained unclear.^12^ Furthermore, these processes are highly dynamic, yet current knowledge is predominantly derived from overexpression and immunolabeling studies, creating significant gaps in our understanding of endogenous protein dynamics. To address these limitations, we used CRISPR-Cas9 technology to generate endogenously GFP-tagged small Rab GTPases associated with recycling pathways. Combined with high-resolution time-lapse imaging of *Salmonella* infection, this approach enabled us to characterize the transient recruitment and spatial organization of recycling pathway components at infection sites.

Our findings support the following integrated model: *Salmonella* effectors SopE and SopE2 that drive the formation of membrane ruffles and infection-associated macropinosomes (IAMs) lead to localized PS accumulation at the infection site. This PS-rich environment serves as a membrane recruitment platform for the deubiquitinase USP32. At the infection site, USP32 deubiquitinates the key recycling regulators Rab14 and Rab11, thereby modulating their effector binding preferences. This PTM-mediated shift in binding specificity redirects Rab14 toward the effector Rufy1 and Rab11 toward FIP3, amplifying recycling dynamics at the infection site. The functional importance of this USP32-dependent pathway is demonstrated by our finding that USP32 silencing substantially impairs Salmonella intracellular replication. Notably, the dependence of *Salmonella* on this host factor identifies USP32 as a promising target for host-directed antimicrobial strategies, particularly given the established drugability of deubiquitinases.^20^

The pivotal roles of Rab11 and Rab14 identified in our study are supported by their known functions in cellular physiology and pathogen-host interactions. Rab11 has been implicated in various cellular stress responses, including those triggered by mechanical or infection-related stimuli, suggesting a role for this GTPase in membrane reorganization in response to stress.^29^ Rab14, in turn, has been shown to regulate cationic substance uptake through endocytic pathways that traffic to non-acidic LAMP1-positive compartments, suggesting a potential function in non-degradative endosomal maturation.^30^ Notably, Rab14 depletion delayed maturation of *Candida albicans*-containing phagosomes to LAMP1-positive compartments,^31^ further supporting its role in this non-degradative trafficking pathway. These observations highlight how *Salmonella* may exploit the inherent properties of these specialized Rabs while providing new insights into their biological functions.

The early steps of *Salmonella* infection are characterized by the formation of long membranous tubules emanating from the SCV.^7,8^ While SVAT formation increases the risk of SCV rupture, concurrent fusion events between the SCV and IAMs replenish and stabilize the SCV membrane.^7^ At present, the precise molecular machinery underlying SVAT formation remains unclear. Both SNX1 and SNX3 have been documented at infection sites,^8,9^ and retromer complex components (specifically Vps35) have been detected at the SCV as early as 15 minutes post-infection,^32^ suggesting that multiple sorting mechanisms may operate simultaneously. This multiplicity is perhaps unsurprising given the substantially larger size of the SCV compared to conventional endosomes, which may facilitate recruitment of diverse sorting machinery at distinct membrane domains. More recently, it was demonstrated that the *Salmonella* effector SopB manipulates ESCRT complex components involved in membrane protein sorting via intraluminal vesicles.^33^ While the functional significance of SopB-ESCRT interactions during *Salmonella* infection remains to be established, this suggests yet another mechanism by which *Salmonella* manipulates host cell biology to control SCV membrane protein composition.

A key unresolved question is which cargo molecules are sorted from the SCV and how they are selectively recognized and directed for removal. We observed CI-M6PR localization along recycling tubules, suggesting selective sorting and removal from the nascent SCV. As CI-M6PR can release its binding to lysosomal enzymes at low pH,^12,34^ CI-M6PR sorting likely represents a mechanism to prevent delivery of lysosomal hydrolases to the SCV lumen. Similarly, a functional SCV must presumably avoid the accumulation of toxic molecules and maintain adequate nutrient availability. Although many transmembrane proteins have been reported to play key roles in *Salmonella* intracellular pathogenesis,^35–38^ the molecular mechanisms governing their selective elimination or retention at the SCV membrane remain largely unexplored. Comprehensively understanding how bacteria manipulate host recycling pathways will provide crucial insights into both the cell-autonomous defense mechanisms that restrict intracellular bacterial replication and the evolutionary counter-strategies that bacteria have developed to overcome or exploit these cellular processes.

## MATERIAL AND METHODS

### Resources Table

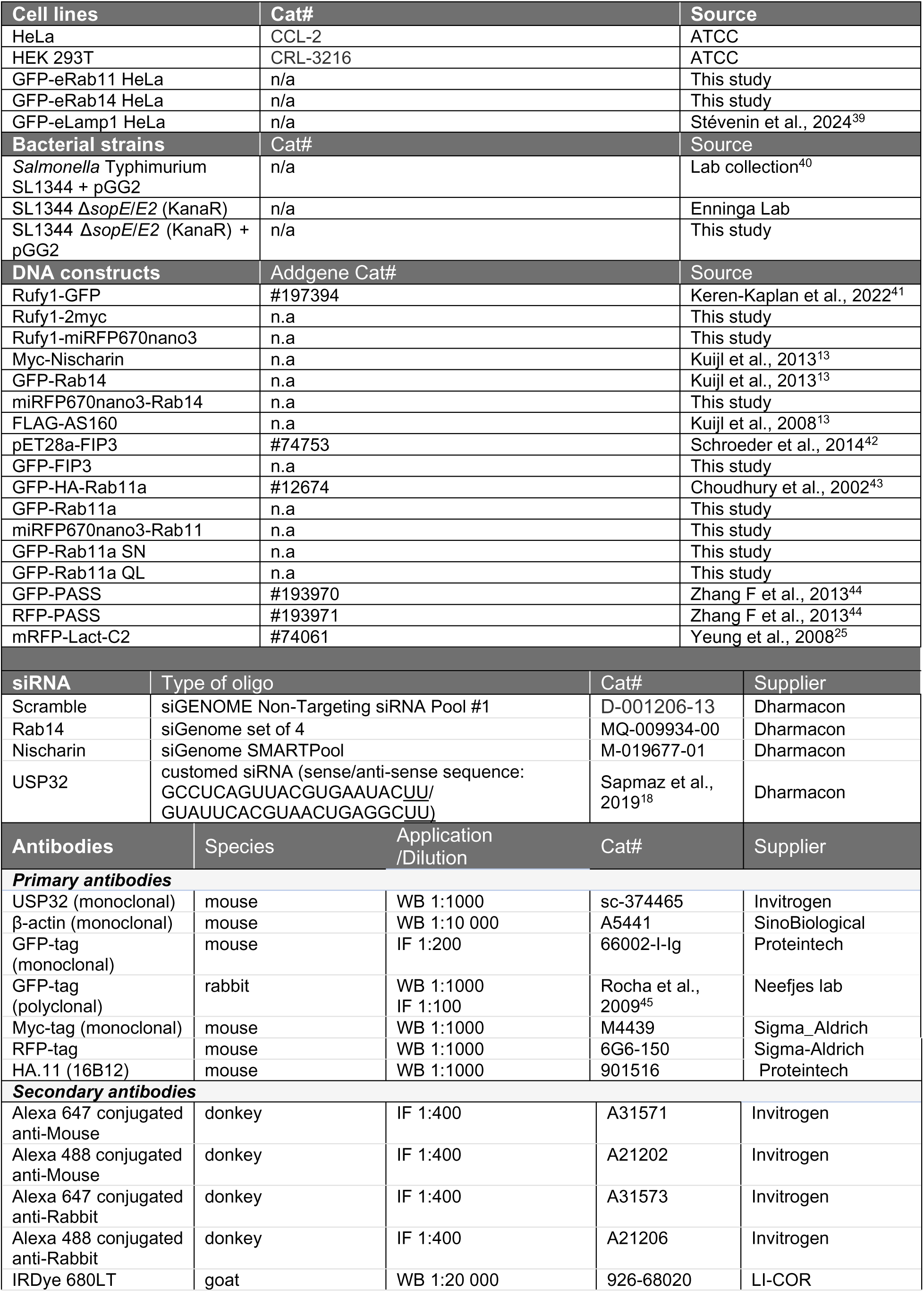
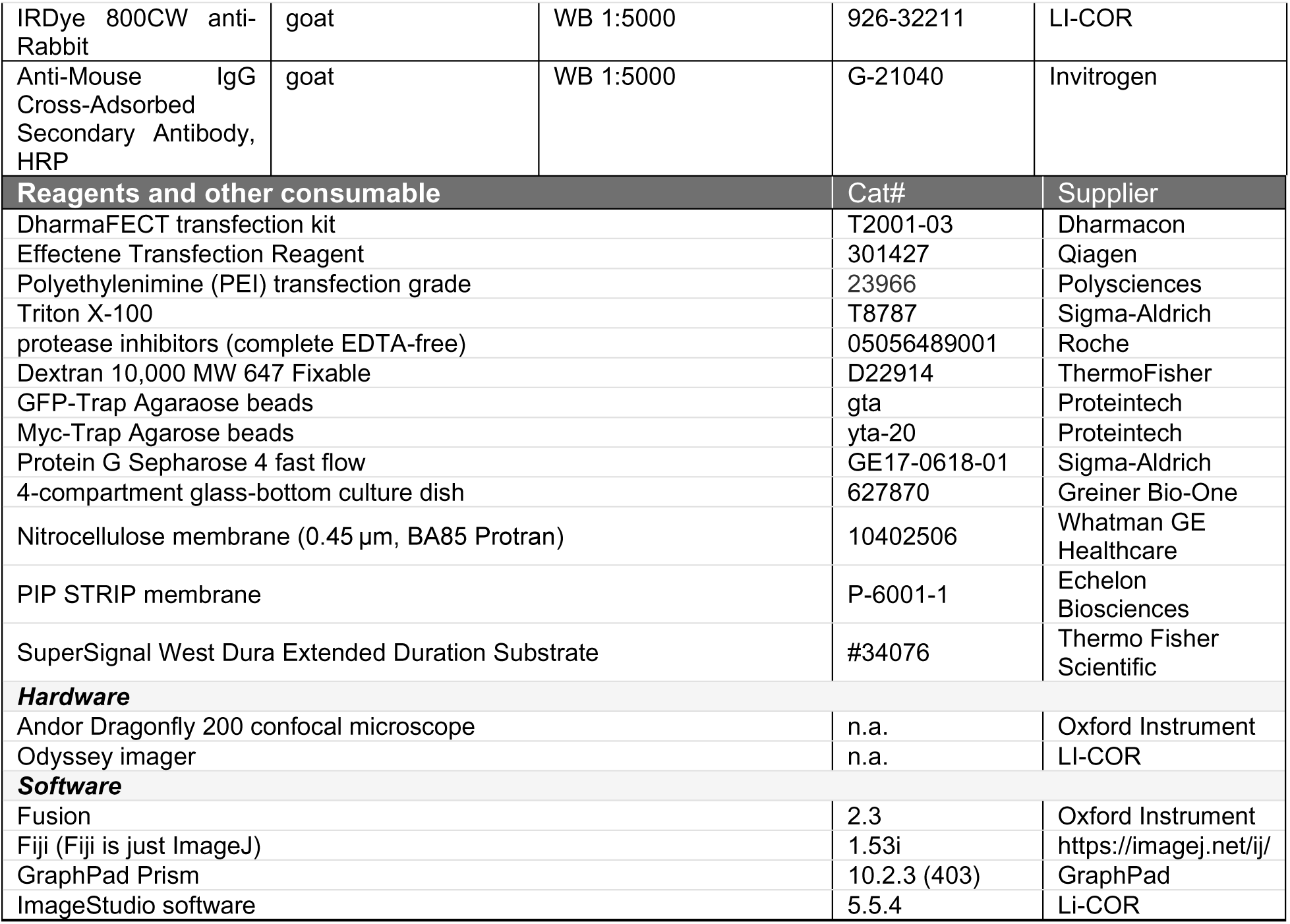

#### Cell culture

HeLa and HEK293T cells were cultured in Dulbecco’s modified Eagle’s medium (DMEM) supplemented with 8% (v/v) Fetal Bovine Serum (FBS). Cell lines were authenticated using short tandem repeat profiling. All cell lines used in the study were cultured at 37°C, 5% CO2, and routinely tested for mycoplasma.

Endogenous Rab11a and Rab14 were tagged with GFP in HeLa cells using CRISPR-CAS9 technology, as described previously^46^. Briefly, a homologous repair construct was made by cloning 250 bp genomic regions up and downstream of the start residing on the first exon from Rab11 and Rab14 which resulted in a mGFP HDR repair construct. gRNAs were designed using the CRISPOR online tool and cloned into pX330 U6 hSpCas9 vector. The primers used are listed in **Table S1.**

Homologous repair and pX330 constructs were co-transfected into HeLa cells using Effectene transfection reagent according to the manufacturer’s instructions. The transfected cells were expanded over 8 days. Next, GFP-positive cells were single-cell sorted with flow cytometry. The single-cell clone-derived cell lines were confirmed through Sanger sequencing and microscopy analyses.

#### Sanger sequencing

Genomic DNA isolates were amplified from cell lines using ISOLATE II Genomic DNA kit (BIO-52066, Bioline) and sequenced using a BigDye Terminator Kit (Applied Biosystems). Sequences were analyzed using Snapgene (GSL Biotech).

#### DNA constructs

miRFP670nano3 (abbreviated here as “670”) is a constitutively fluorescent protein with low acid sensitivity (excitation/emission wavelengths: 647/670 nm).^47^ Rufy1 fusion proteins were generated as follows. Rufy1-GFP was digested with XhoI and Asp718I to excise the Rufy1 fragment, which was then cloned into the vectors 2Myc-N1 and miRFP670nano3-N1 to generate Rufy1-myc and Rufy1-670, respectively. GFP-FIP3 was generated by PCR amplification and cloned into mGFP-C1 (Clontech) using BglII and EcoRI restriction sites. 670-Rab11, 670-Rab14, and 670-FIP3 were generated by swapping mGFP for miRFP670nano3 in the corresponding mGFP-tagged constructs (mGFP-Rab11, mGFP-Rab14, and mGFP-FIP3). All primer sequences are provided in Table S1.

#### siRNA and DNA transfection

For siRNA transfection, cells were diluted at a 1:3 ratio 24 h prior to transfection to optimize transfection efficiency. siRNA transfections were performed using the DharmaFECT transfection kit according to the manufacturer’s instructions, 72 h before infection, imaging, or cell lysis. For DNA transfection, cells were transfected for 16 h using Effectene Transfection Reagent (for microscopy) or PEI transfection reagent (for co-immunoprecipitation), according to the respective manufacturer’s instructions. In experiments combining siRNA and DNA transfection, the medium was replaced before DNA transfection to reduce cytotoxicity. All reagents, constructs, and siRNA sequences are listed in the Resources tables.

#### Bacterial culture

*Salmonella enterica* serovar Typhimurium strains SL1344 (wild-type and ΔsopE/E2) were used for all infection experiments. Bacteria were transformed with the pGG2 plasmid (rpsm::dsRed) to enable constitutive expression of dsRed.^48^ Bacterial cultures were grown in Lysogeny Broth (LB) medium supplemented with ampicillin (50 μg/mL) for plasmid selection and maintenance.

#### Bacterial infection

The day before the infection, bacteria were cultured overnight in 5 mL LB medium at 37°C with orbital shaking (220 rpm). Growth medium was supplemented with 50 μg/mL ampicillin for plasmid selection. On the day of the infection, bacterial subcultures were prepared by diluting the overnight culture 1:33 into fresh LB supplemented with antibiotics and incubated at 37°C with orbital shaking (220 rpm) for 3 h until late exponential phase. Then, 1 mL of bacteria subculture was centrifuged at 9,000 *g* for 1 min. The supernatant was discarded, and the pellet was resuspended in EM buffer (120 mM NaCl, 7 mM KCl, 1.8 mM CaCl_2_, 0.8 mM Mg Cl_2_, 5 mM glucose, 25 mM HEPES, pH 7.3). This centrifugation and resuspension step was repeated once to wash the bacteria. Bacteria were then diluted in EM buffer to a multiplicity of infection (MOI) of 20 to prepare the infection mix. The cell medium was replaced by 250 μL of infection mix. For live-cell imaging of bacterial entry, cells were immediately transferred to the microscope for field selection. For dextran labeling experiments, the infection mix was supplemented with Dextran (10,000 MW, Alexa Fluor 647 conjugate), and cells were incubated with bacteria for 30 min at 37°C with 5% CO_2_. Cells were then washed three times with warm EM buffer to remove Dextran and extracellular bacteria. The medium was replaced with EM buffer supplemented with 10% fetal bovine serum (FBS) and gentamicin (20 μg/mL). Time-lapse microscopy was initiated immediately after the washing steps (defined as t=W+0).

#### Fixed microscopy

Cells were seeded onto glass coverslips (thickness No. 1.5) in 24-well plates. At the desired time point post-infection, cells were fixed with 4% paraformaldehyde for 15 min in the dark at room temperature (RT). Fixed cells were washed three times with PBS and stored at 4°C for a maximum of one week. Before staining, samples were blocked in PBS supplemented with 20% fetal bovine serum (FBS) and 0.25% saponin (for gentle permeabilization) for 1 h at RT in the dark. Samples were immunolabeled with primary antibody for 1 h at RT in the dark, washed with PBS, and incubated with secondary antibodies and DAPI for 45 min at RT in the dark. Samples were washed again in PBS and mounted in Prolong Gold. Antibody dilutions and references are provided in the Resources tables. Confocal microscopy was performed on an Andor Dragonfly 200 spinning-disk microscope equipped with a 63×/1.40–0.60 oil Plan Apo objective, and an Andor Zyla 4.2 PLUS sCMOS camera.

#### Time-lapse microscopy

Cells were seeded into a four-compartment glass-bottom culture dish days prior to the infection (72 h for siRNA-transfected cells, 48-24 h for DNA-transfected or non-transfected cells). Following cell infection under a laminar flow hood, the dish was transferred to a 37°C, 5% CO₂ humidified chamber mounted on an Andor Dragonfly 200 spinning-disk microscope. Three to five fields of view per condition were selected and imaged using a 40×/1.30 oil HCX Plan Apo objective or a 63×/1.40–0.60 oil Plan Apo objective. For each field of view, a Z-stack spanning 5.5 μm (0.3 μm step size) was acquired in each channel. Total duration and frequency of acquisitions were adapted depending on the number of channels and

#### Microscopy acquisition display

Unless otherwise specified in the figure caption, images display maximum intensity Z-projections generated using the image analysis software Fiji. Cyan, Red, and Yellow lookup tables (LUTs) were used for overlay images to ensure accessibility for all readers, including those with color-blindness. The Inferno LUT was used to highlight signal intensity variations.

Time-lapse series display time points selected to highlight the most relevant events; these are not uniformly spaced in time. Complete infection events are shown in Supplementary Videos.

#### Image analysis

Quantification of time-lapse data was performed manually using Fiji in a blinded manner. The fraction of Rab14-positive SCVs was determined by counting the number of SCVs that recruited Rab14 during the 2 h acquisition period, divided by the total number of SCVs analyzed. Similarly, the fraction of Rab11-positive infection sites was determined by counting the number of infection sites that recruited Rab11 during the 2 h acquisition period, divided by the total number of infection sites analyzed. GFP-Rab11 recruitment intensity was quantified by measuring the raw integrated density of the GFP signal using the Fiji “Measure → Raw Integrated Density” function within a circular region of interest (147 pixels in diameter) centered at the infection site for each time point. The recruitment amplitude was calculated as the difference between the maximum intensity and the intensity at t = 0. Intensity versus time graphs were generated for each infection site to enable manual verification and artifact detection.

#### SDS-PAGE and Western blot

The proteins of the pull-down and whole-cell lysate were separated by a 10% SDS-PAGE and transferred to a nitrocellulose membrane at 300 mA for 1.5 h. Membranes were blocked in 5% milk in PBS. Primary antibodies were diluted in 5% milk in 0.1% PBS-Tween 20 (PBST). Membranes were incubated at RT with antibody solutions for 1h under constant agitation, washed 3 times for 5 min in PBST, incubated with fluorogenic secondary antibodies in 5% milk in 0,1% PBST for 45-60 min, and washed 3 times for 5 min in PBST. The fluorescent signal was recorded with the Odyssey imaging system and Image Studio software (Li-Cor). Band intensity was quantified using ImageJ or Image studio Software.

#### Ubiquitination assay

HEK293T cells were transfected with HA-ubiquitin and other constructs of interest for 24 h, collected by scraping in 300 µL lysis buffer 1 (50 mM Tris-HCl, pH 7.5, 150 mM NaCl, 5 mM EDTA, 0.5% Triton X-100, 10 mM N-methyl maleimide, and EDTA-free protease inhibitors cocktail). Then, 100 µL of lysis buffer 2 (100 mM Tris-HCl, pH 8.0, 1 mM EDTA, and 2% SDS) was added to the lysates, and the samples were sonicated. Next, the SDS was diluted by raising the volume of the samples to 1 mL with lysis buffer 1. The samples were then centrifuged at 20,000 *g* for 20 min at 4 °C. Cell lysates were incubated overnight at 4 °C with 6 µL GFP Trap beads. Afterward, the beads were washed 3 times with lysis buffer. Then, 30 µL Protein G Sepharose 4 fast flow was added in a fourth wash. All the liquid was removed from the beads before SDS sample buffer was added (supplemented with 10 mM DTT). The proteins were denatured at 95 °C for 15 min and separated on 8% SDS-PAGE gel.

#### Co-immunoprecipitation

HEK293T cells were lysed in lysis buffer containing 50 mM Tris-HCl at pH 7.5, 50 mM NaCl, 0.8% NP-40, 5mM MgCl_2,_ and protease inhibitors (Roche Diagnostics, EDTA-free) and incubated for 30 min under gentle agitation at 4°C. Supernatant after spinning (10min at 12,000 *g* at 4°C) was incubated with 8 µL myc-trap or GFP-trap beads (Chomotek) for 2 h at 4°C. Part of the supernatant was kept as whole-cell lysate (WCL). Beads were washed four times in lysis buffer by centrifugating the samples (1 min., 4°C, 500 *g*) before the addition of Laemmli Sample Buffer (containing 5% β-mercaptoethanol), followed by 10 min incubation at 96°C. CoIP proteins were separated by SDS-PAGE for Western blotting and detection by antibody staining.

#### Protein Expression and Purification

Full-length USP32 (amino acids 1-1604) was expressed in *Spodoptera frugiperda* Sf9 insect cells and purified as previously described.^18^

#### Phospholipid Binding Assay

USP32 phospholipid binding was assessed using PIP Strip membranes (P-6001-1, Echelon Biosciences). Membranes were blocked with 5 mL PBS-T (PBS + 0.1% Tween-20) containing 3% bovine serum albumin (BSA) for 1 h at RT with gentle agitation. Membranes were then incubated with 2 μg/mL purified USP32 protein diluted in 5 mL PBS-T + 3% BSA for 1 h at RT with gentle agitation. Following three 5-min washes with PBS-T, membranes were incubated with mouse anti-USP32 antibody (1:1000 in PBS-T + 3% BSA) for 1 h at RT with gentle agitation. After three additional 5-min PBS-T washes, membranes were incubated with HRP-conjugated anti-mouse secondary antibody (1:5000 in PBS-T + 3% BSA) for 1 h at RT with gentle agitation, followed by three final 5-min PBS-T washes. Bound protein was detected using enhanced chemiluminescence (SuperSignal West Dura Extended Duration Substrate) according to the manufacturer’s instructions. Chemiluminescent signals were captured using an AI600 imaging system (GE Healthcare) with automated exposure settings.

#### Quantification and Statistical Analysis

Statistical analyses of microscopic acquisitions or Western blots were performed using GraphPad Prism. t-tests or ordinary one-way ANOVA were used to evaluate the significance of the results, referred to as *, **, ***, and **** for p values < 0.05, < 0.01, < 0.001, and < 0.0001, respectively. Statistical details of experiments can be found in the figure legends.

## Acknowledgments

This work was supported by a Veni grant (VI.Veni.222.124) from the Dutch Research Council (NWO) to VS and a Spinoza award of the Dutch Research Council (“Nederlandse Organisatie voor Wetenschappelijk Onderzoek”; NWO 00897590) awarded to JN. JN is an Oncode Institute investigator. AS is funded by the Institute for Chemical Immunology (grant no. ICI00026) and the Innovative Medicines Initiative 2 (IMI2) Joint Undertaking under grant agreement no. 875510 (EUbOPEN project). The effectors-deleted Salmonella strains were kindly provided by Dr. Valenzuela Montenegro and Dr. Enninga. The authors thank all the kind Addgene plasmid depositors (see Resources Table and references).

## Author contributions

A.S. and V.S. designed the project. A.S., S.Y.v.d.Z, L.M., E.F., N.H., and V.S. performed the experiments and analyses. J.J.L.L.A. and L.J. produced cell lines and constructs. J.N. provided infrastructure and funding. J.N. and I.B. provided scientific feedback. V.S. supervised the study and wrote the manuscript. All authors read, edited, and approved the manuscript.

## Declaration of interests

The authors declare no competing interests.

## SUPPLEMENTARY DATA

Supplementary Videos and Table access: https://drive.google.com/drive/folders/10DjUdyOjEhy-IswvUkpbY2rys42a6Bug?usp=sharing

**Video S1.** Time-lapse microscopy of HeLa cells overexpressing GFP-SNX1 (yellow), and 670-Rab14 (magenta) and infected with dsRed-expressing *Salmonella* (cyan). Bacteria are added to the cells just before the beginning of the acquisition to visualize entry and early events. Scale bars: 5 µm. Related to Fig. 1A.

**Video S2.** Time-lapse microscopy of HeLa cells overexpressing GFP-SNX1 (yellow), and 670-Rab11 (magenta) and infected with dsRed-expressing *Salmonella* SL1344 WT (cyan). Bacteria are added to the cells just before the beginning of the acquisition to visualize entry and early events. Scale bars: 5 µm. Related to Fig. 1B.

**Video S3.** Time-lapse microscopy of HeLa cells endogenously GFP-tagged for Rab14 (Inferno LUT) and infected with dsRed-expressing *Salmonella* (cyan). Bacteria are added to the cells just before the beginning of the acquisition to visualize entry and early events. Scale bars: 5 µm. Related to Fig. 2A.

**Video S4.** Time-lapse microscopy of HeLa cells endogenously GFP-tagged for Rab11 (Inferno LUT) and infected with dsRed-expressing *Salmonella* (cyan). Bacteria are added to the cells just before the beginning of the acquisition to visualize entry and early events. Scale bars: 5 µm. Related to Fig. 2B.

**Video S5.** Time-lapse microscopy of HeLa cells endogenously GFP-tagged for Rab14 (magenta) and infected with dsRed-expressing *Salmonella* (cyan) in the presence of Dextran (yellow). Dextran and bacteria are added to the cells for 30 min and washed before the beginning of the acquisition to visualize IAM dynamics. Scale bars: 5 µm. Related to Fig. 2E.

**Video S6.** Time-lapse microscopy of HeLa cells endogenously GFP-tagged for Rab11 (yellow), overexpressing 670-Rab14 (magenta) and infected with dsRed-expressing *Salmonella* (cyan). Bacteria are added to the cells just before the beginning of the acquisition to visualize entry and early events. Scale bars: 5 µm. Related to Fig. 2K.

**Video S7.** Time-lapse microscopy of HeLa cells endogenously GFP-tagged for Rab14 (yellow), overexpressing 670-Rab11 (magenta) and infected with dsRed-expressing *Salmonella* (cyan). Bacteria are added to the cells just before the beginning of the acquisition to visualize entry and early events. Scale bars: 5 µm. Related to Fig. S2.

**Video S8.** Time-lapse microscopy of HeLa cells endogenously GFP-tagged for LAMP1 (magenta) and infected with dsRed-expressing *Salmonella* (cyan) under siCtrl (left) versus siUSP32 (right) conditions. Bacteria are added to the cells for 30 min and washed before the beginning of the acquisition. Scale bars: 50 µm. Related to Fig. 3F.

**Video S9.** Time-lapse microscopy of HeLa cells overexpressing Rufy1-GFP (Inferno LUT) and infected with dsRed-expressing *Salmonella* SL1344 WT (cyan). Bacteria are added to the cells just before the beginning of the acquisition to visualize entry and early events. Scale bars: 5 µm. Related to Fig. 4G.

**Video S10.** Time-lapse microscopy of HeLa cells overexpressing GFP-FIP3 (Inferno LUT) and infected with dsRed-expressing *Salmonella* SL1344 WT (cyan). Bacteria are added to the cells just before the beginning of the acquisition to visualize entry and early events. Scale bars: 5 µm. Related to Fig. 5C.

**Table S1.** List of primers used in the study.

**Fig. S1.**
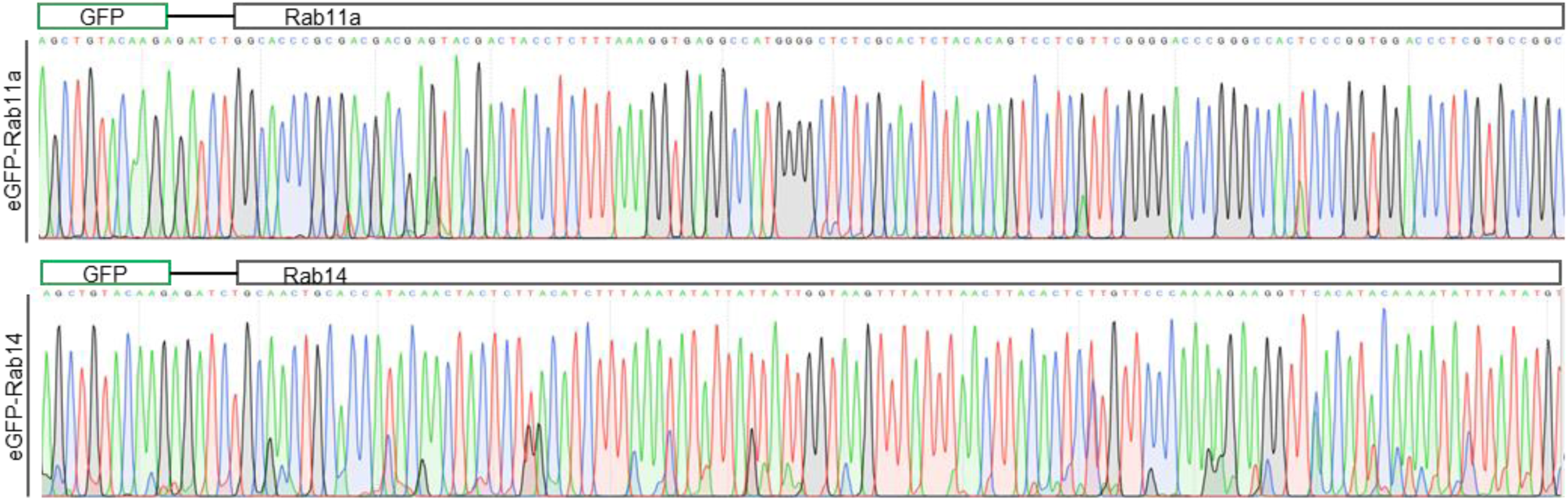
Sanger sequencing data of clonally derived endogenous GFP-tagged Rab11a and Rab14 HeLa cell lines.

**Fig. S2.**
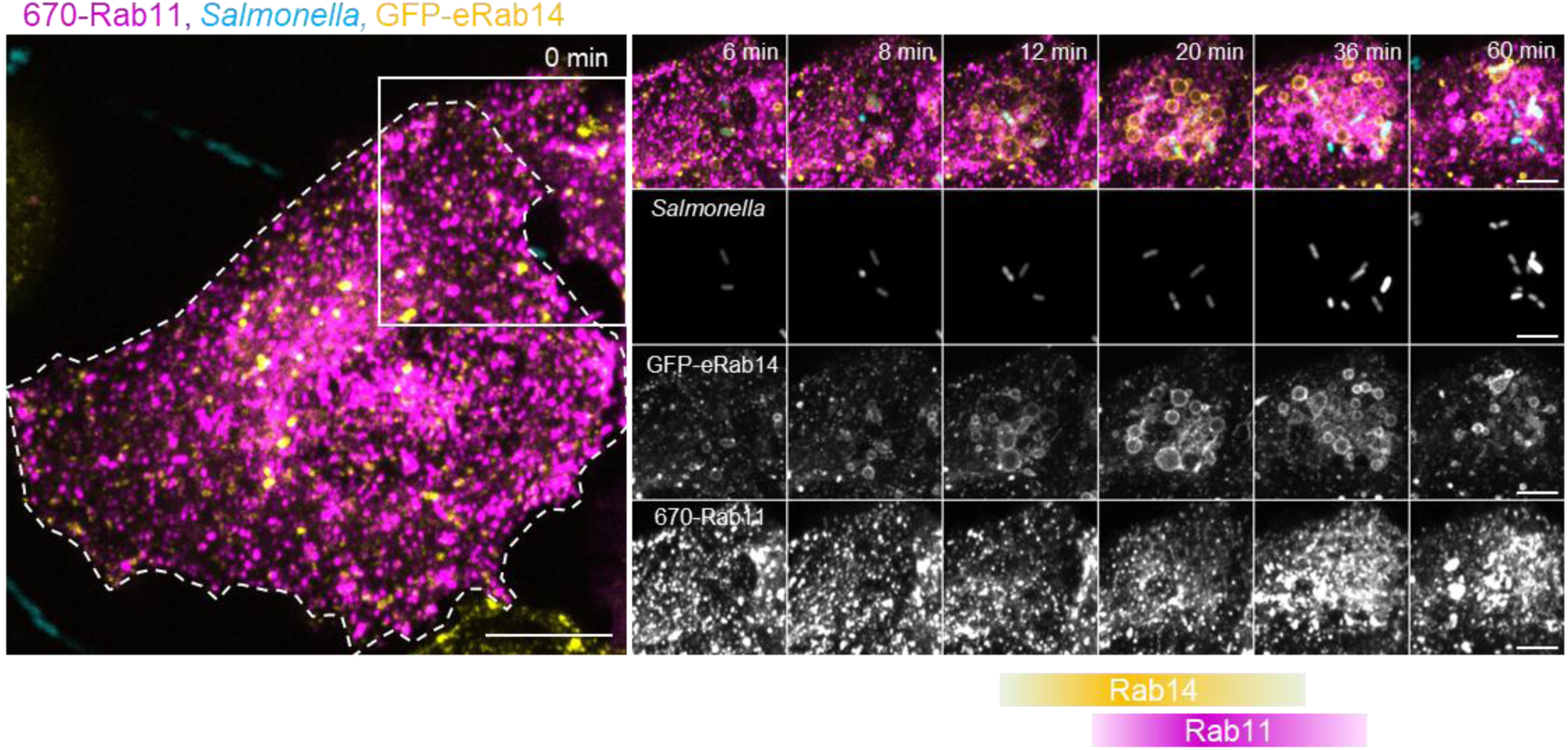
Time-lapse microscopy of HeLa cells endogenously GFP-tagged for Rab14 (yellow), overexpressing 670-Rab11 (magenta), and infected with dsRed-expressing *Salmonella* (cyan). Bacteria were added immediately before acquisition to capture bacterial entry and early recruitment events. See Video S7. Scale bars: 10 μm (overview) and 5 μm (inset).

**Fig. S3.**
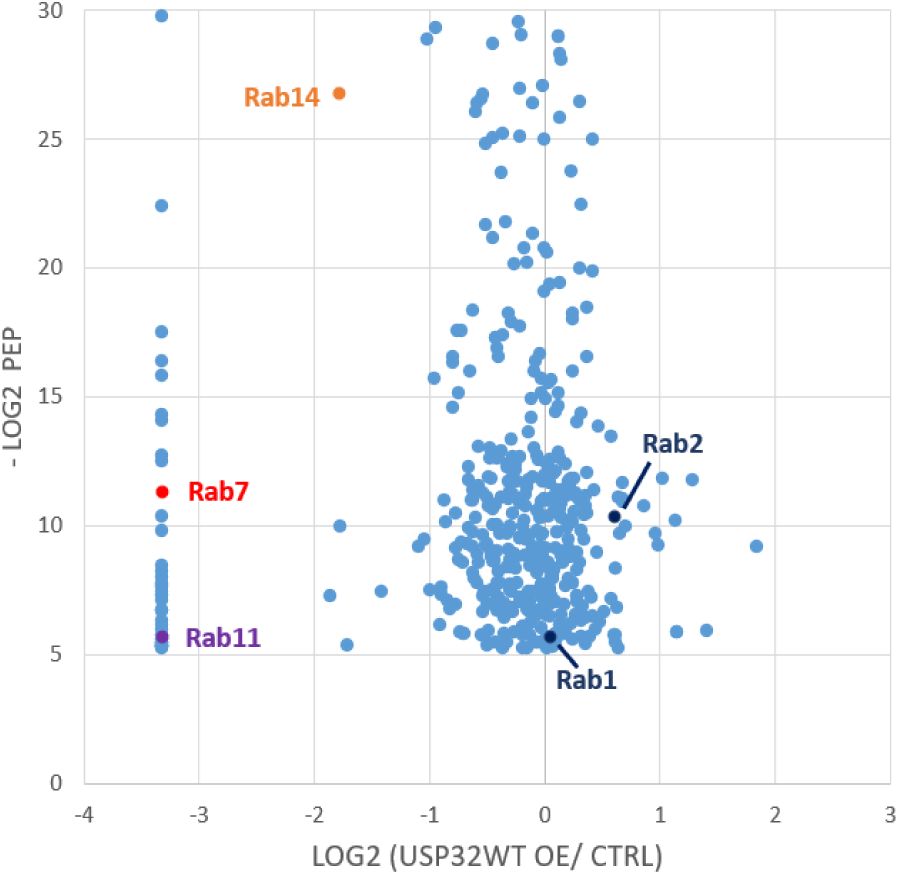
HEK cell ubiquitome upon USP32-HA overexpression versus vector control (Ctrl), based on a published mass spectrometry dataset,^18^ accessed via the PRIDE repository PXD011899. All identified Rabs are labeled in the volcano plot.

**Fig. S4.**
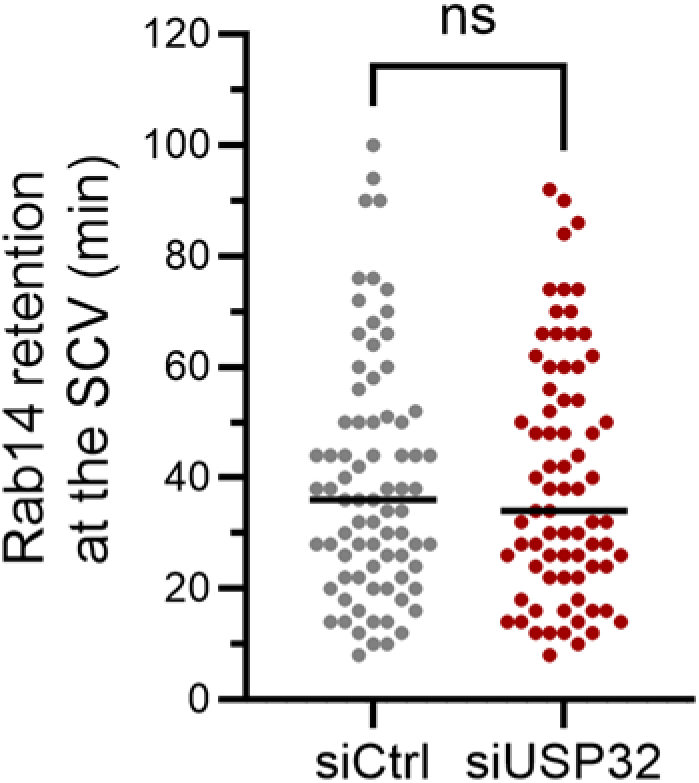
Quantification of Rab14 retention time at SCVs monitored by time-lapse microscopy over 2 h of infection in siCtrl versus siUSP32 conditions. Each point represents one SCV (n = 3 biological replicates). Statistical analysis by Student’s t-test; ns, p > 0.05.

**Fig. S5.**
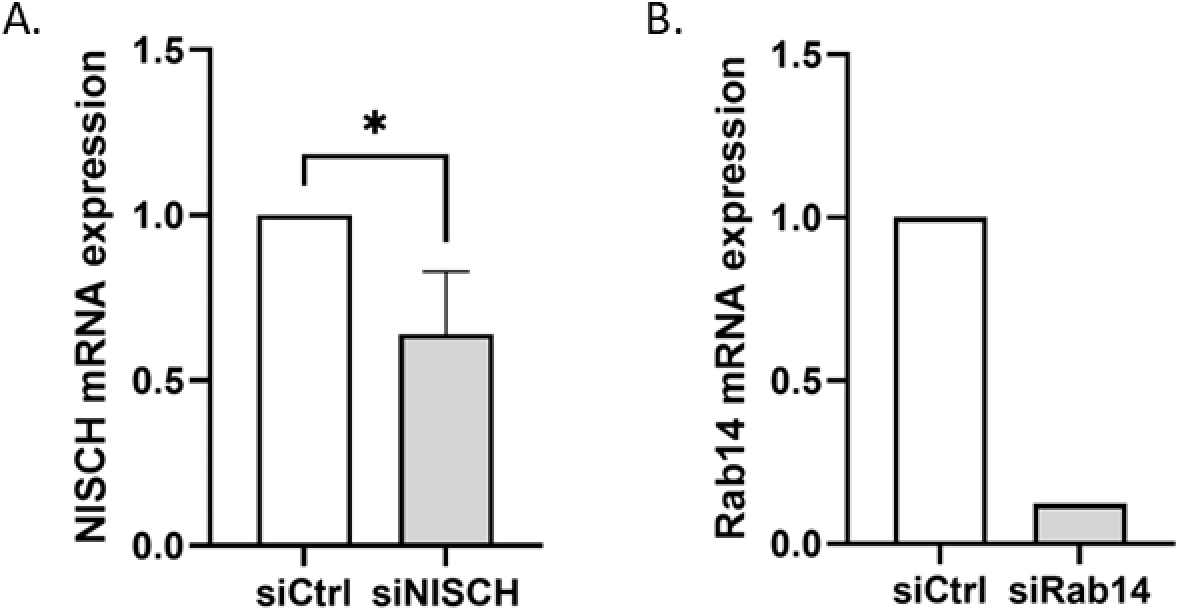
RT-qPCR validation of Nischarin (A) and Rab14 (B) silencing after 72h of siRNA treatment.

**Fig. S6.**
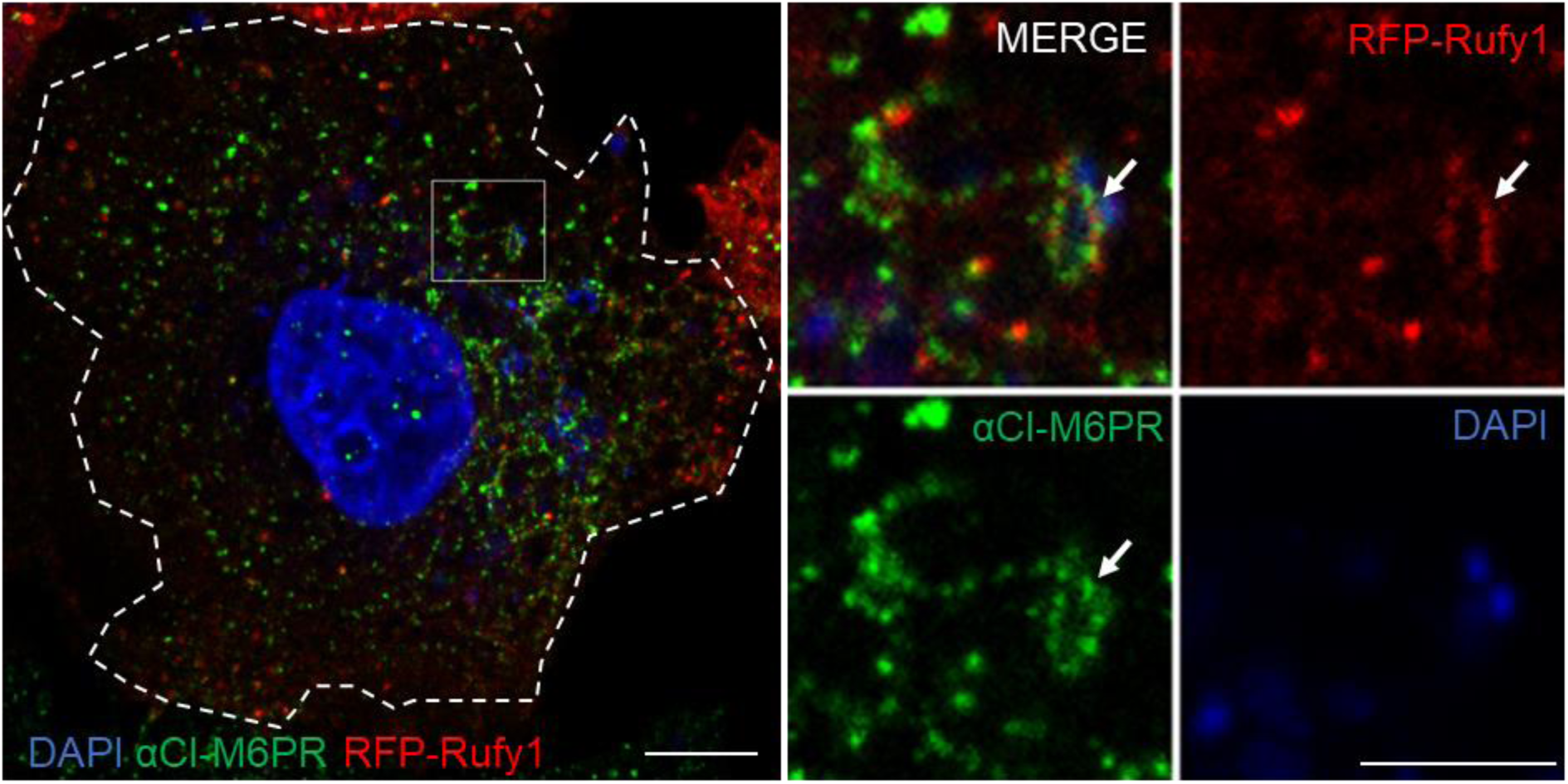
Immunofluorescent detection of CI-M6PR (green) in HeLa cells overexpressing RFP-Rufy1 (red), infected with *Salmonella* and fixed at 1 hpi. Scale bars: 10 μm (overview) and 5 μm (inset). A single z-plan is displayed.

**Fig. S7.**
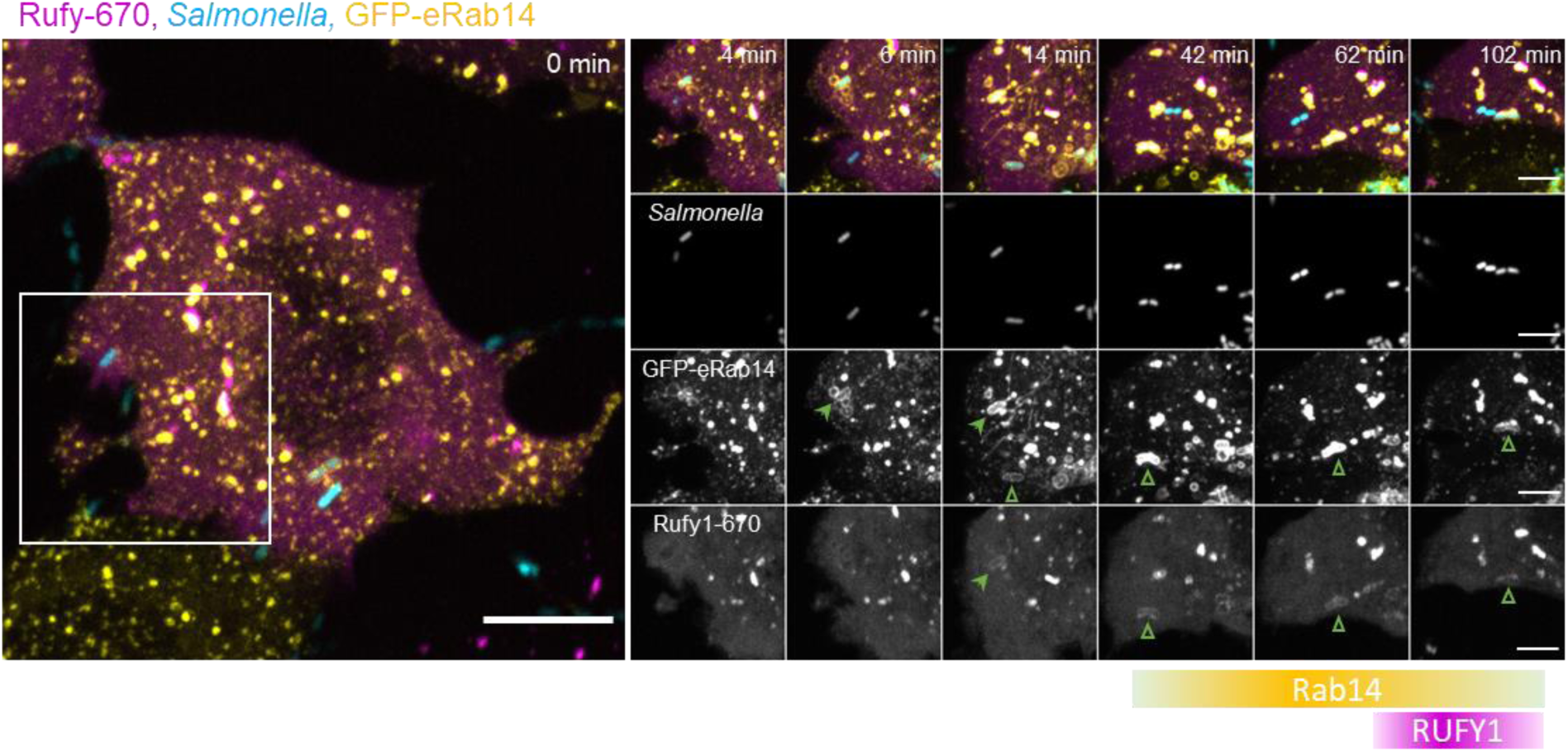
Time-lapse microscopy of HeLa cells endogenously GFP-tagged for Rab14 (yellow), overexpressing Rufy1-670 (magenta), and infected with dsRed-expressing *Salmonella* (cyan). Bacteria were added immediately before acquisition to capture bacterial entry and early recruitment events. Scale bars: 10 μm (overview) and 5 μm (inset). Representative of n = 3 biological replicates. Green arrows indicate SCVs, highlighting the sequential recruitment of Rab14 and Rufy1.

**Fig. S8.**
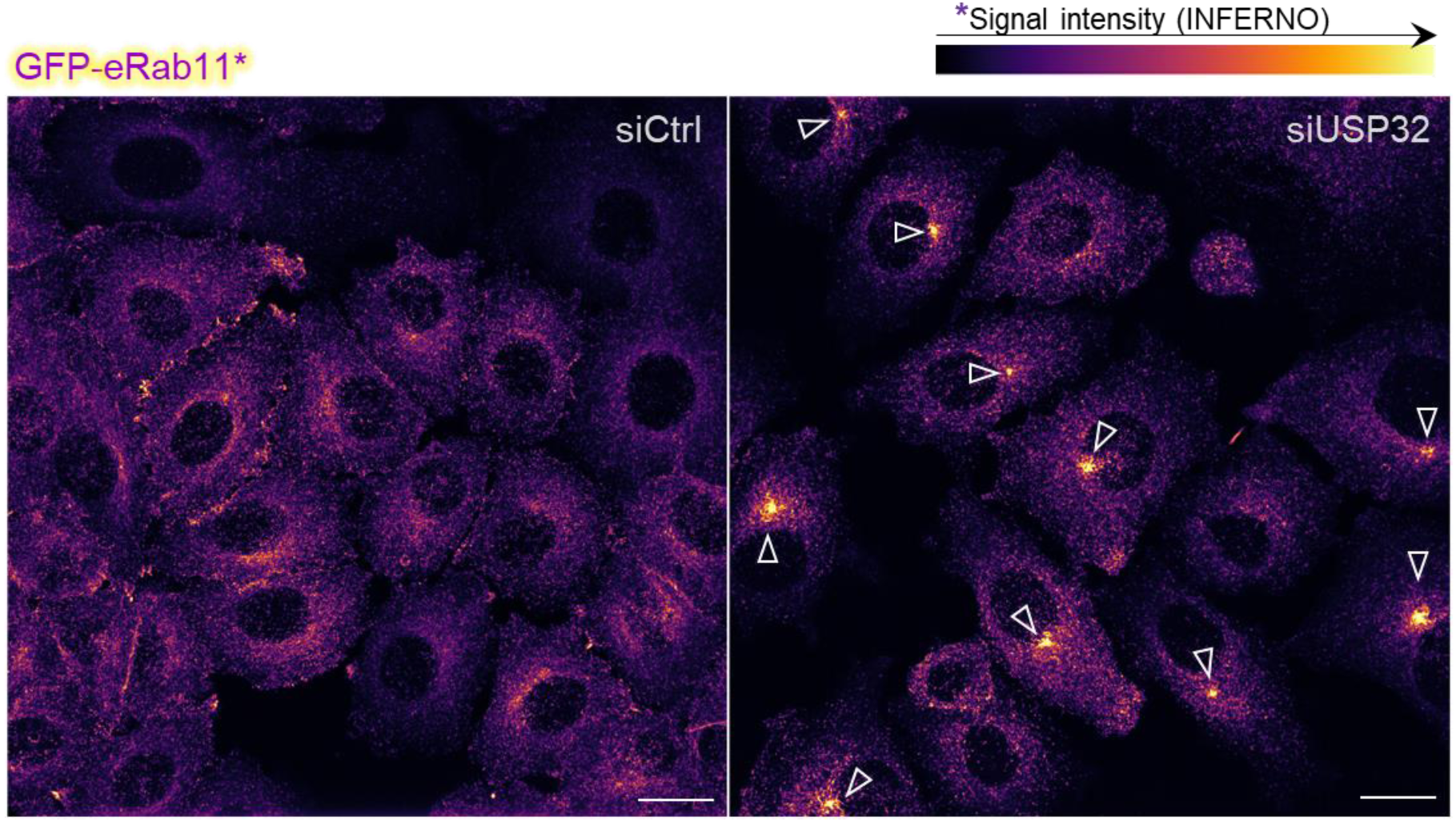
Representative microscopy image of HeLa cells endogenously GFP-tagged for Rab11 (Inferno LUT) in siCtrl and siUSP32 conditions. Arrows indicate perinuclear accumulations of GFP-eRab11, likely corresponding to the centrosome. Scale bar: 20 µm.

**Fig. S9.**
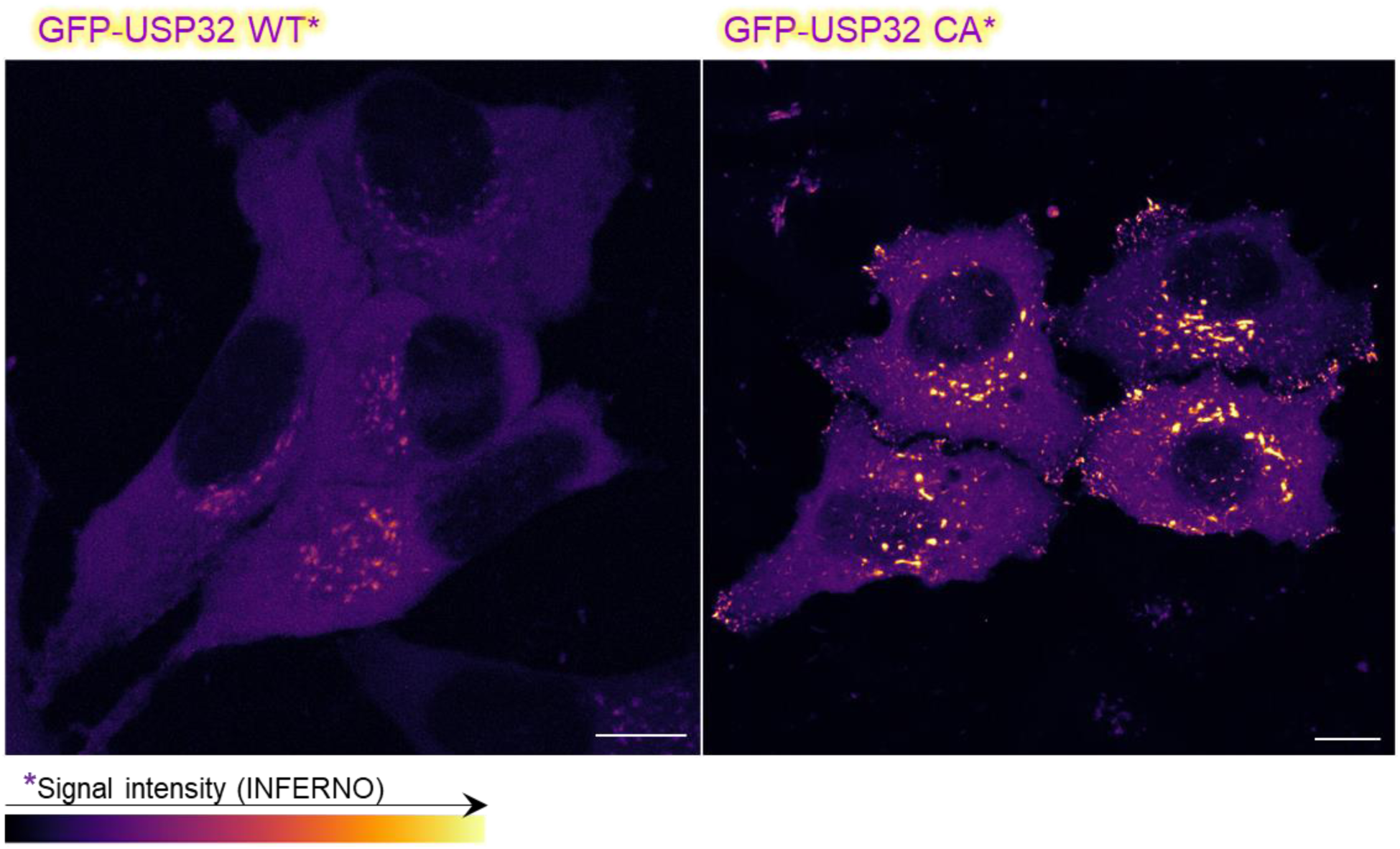
Live cell imaging of HeLa cells overexpressing GFP-USP32 WT or CA (Inferno LUT). Scale bar: 10 µm.

## Notes

https://drive.google.com/drive/folders/10DjUdyOjEhy-IswvUkpbY2rys42a6Bug?usp=sharing

### Competing Interest Statement

The authors have declared no competing interest.

